# Direct high-throughput deconvolution of non-canonical bases via nanopore sequencing and bootstrapped learning

**DOI:** 10.1101/2024.12.02.625113

**Authors:** Mauricio Perez, Michiko Kimoto, Priscilla Rajakumar, Chayaporn Suphavilai, Rafael Peres da Silva, Hui Pen Tan, Nicholas Ting Xun Ong, Hannah Nicholas, Ichiro Hirao, Chew Wei Leong, Niranjan Nagarajan

## Abstract

The discovery of non-canonical bases (NCBs) in viruses and the development of synthetic xeno-nucleic acids (XNAs) to expand the genetic alphabet has spawned interest in many applications, from viral genomics, to synthetic biology and DNA storage. However, the inability to read non-canonical bases in a direct, high-throughput manner has been a significant limitation to its study and applicability. Here we demonstrate that XNA templates containing non-canonical bases can be directly and robustly sequenced (>2.3 million reads/flowcell, similar to DNA controls) on a MinION sequencer from Oxford Nanopore Technologies to obtain signal data that is significantly distinct from DNA controls (median fold-change >6×). To enable training of machine learning models that deconvolve these signals and basecall non-canonical and canonical bases, we developed a framework to synthesize a complex pool of 1,024 NCB-containing oligonucleotides with diverse 6-mer sequence contexts and high purity (>90% NCB-insertion on average). Bootstrapped models to assist in data preparation, and data augmentation with spliced reads to provide high context diversity, enabled learning of a generalizable model to call canonical as well as non-canonical bases with high accuracy (>80%) and specificity (99%). These results highlight the versatility of nanopore sequencing as a platform for interrogating nucleic acids for viral genomic and xenobiology applications, and the potential to transform the study of genetic material beyond those that use canonical bases.

## Introduction

Synthetic xeno-nucleic acids (XNAs) consist of unnatural bases (UBs) beyond the canonical A, T, C, G, and U found in naturally occurring DNA and RNA. Such non-canonical bases (NCBs) expand the information space of the genetic code, offer novel biochemical properties, and have opened up new fields in xenobiology, synthetic biology, and biotechnology. Exemplary XNAs include the bases isoG and isoC developed at the Swiss Federal Institute of Technology^1^, P and Z developed at the Foundation for Applied Molecular Evolution^2,3^, d5SICS and dNaM developed at the Scripps Research Institute^4,5^, Ds and Pa/Px developed at RIKEN^6,7^, and 7-Deaza-2′-deoxyisoguanosine developed at the Center for Nanotechnology^8^. There are also naturally occurring NCBs which have been found in bacterial tRNAs^9^ or predicted to be part of the genomes of diverse viruses as a defense against host restriction systems^9–12^, though direct detection and characterization of their function has been hampered by the lack of appropriate sequencing technologies. The use of NCBs to expand the genetic alphabet has several applications in synthetic biology, including the development of sensors, aptamers, nanostructures, and semi-synthetic organisms^13–15^. With the advent of DNA storage as a paradigm to develop low-energy, ultra-high-density systems^16,17^, reading and writing NCBs has another potential application, however current approaches are limited in speed, nucleotide-resolution, and throughput.

A common early approach to detect NCBs (focusing on UBs) was to use Sanger sequencing and decipher UB positions through signal gaps stemming from the inability to sequence through these UBs^6,7,18,19^. An alternative approach has involved using PCR to replace UBs followed by sequencing and mutation analysis to detect them, but with the drawback of introducing unintended errors during amplification^2,20^. Both approaches suffer from the corresponding limitations of sample processing, which introduce sources of error and operational cost. Next-generation sequencing, while much less limited in throughput, largely faces similar challenges due to its inability to natively detect UBs^21,22^. In particular, the detection of UBs in specific sequence contexts or in the presence of multiple UBs can be challenging^20,22^. A potential alternative is direct single-molecule interrogation of nucleic acids with third-generation technology using nanopores, which have shown promise with canonical nucleic acid base modifications (e.g. methylated DNA^23–26^), though the ability to directly sequence XNAs, which contain a different nucleic acid backbone, has not been demonstrated^27^. Although PacBio is also a third-generation sequencing platform, its sequencing-by-synthesis process may need major adaptations with fluorescently labeled non-canonical bases to facilitate direct sequencing of XNAs.

A major obstacle for direct single-molecule sequencing of non-canonical bases is the ability to synthesize and utilize a sufficiently complex dataset for training basecalling models. In this study, we have addressed these challenges and developed technology that enables direct sequencing of XNAs using a widely available nanopore sequencing platform (Oxford Nanopore Technologies – ONT). Our results suggest that NCB-containing XNA templates can be robustly sequenced on an ONT MinION system, with yield similar to DNA controls (>2 million reads per flowcell), without notable truncation of reads, and with raw signals that are significantly different from canonical bases (median fold-change >6×). To enable direct decoding of XNAs from these raw nanopore signals, we developed an XNA design and synthesis framework that generates a complex library of XNA oligonucleotides (n=1,024) containing all possible single-UB, 6-mer sequences, which provides the critical materials to train a basecaller model for XNAs. We have expanded the conventional basecaller architecture decoding states for it to be able to learn how to also decode the non-canonical bases X and Y, in addition to the canonical bases (A, T, C and G). Via our bootstrapped training approach, which retrieves additional accurate training data, and our read-splicing based data-augmentation technique, responsible for generating reads with high sequence context diversity, we show that a generalizable deep learning model based on convolutional neural networks (CNNs) can be trained to call non-canonical bases with high accuracy (>80%) and specificity (99%). Our work provides the first example of direct high-throughput deconvolution of non-canonical bases using nanopore sequencing, enabling determination of nucleotide-level identities in XNAs needed for synthetic biology applications, and opening up the possibility that other classes of synthetic and natural nucleic acids may also be amenable to a bootstrapped and data-augmented learning approach.

## Results

### Analysis of XNAs on a high-throughput nanopore sequencer retains high fidelity

As a proof-of-concept to study the sequencing of NCB-containing templates, we leveraged the Px-Ds (X-Y) architecture, one of the three known classes of unnatural basepairs^7^ (**Figure 1A**, **Supplementary Figure 1**). The Px-Ds basepair system was designed taking into consideration hydrophobicity, shape, and electrostatic complementarity properties that enable the basepair to be efficiently amplified with PCR^7^, facilitating synthesis of XNAs with NCBs in diverse sequence contexts. For the initial experiment, we designed 20 individual templates with either a single NCB (XNA01-XNA12), multiple NCBs at varying distances (XNA13-XNA16), or four equidistant NCBs (XNA17-XNA20), flanked by stretches of canonical bases in different combinations and tagged via unique 24-nucleotide barcodes (**Supplementary Figure 2**, **Supplementary File 1**).

**Figure 1.**
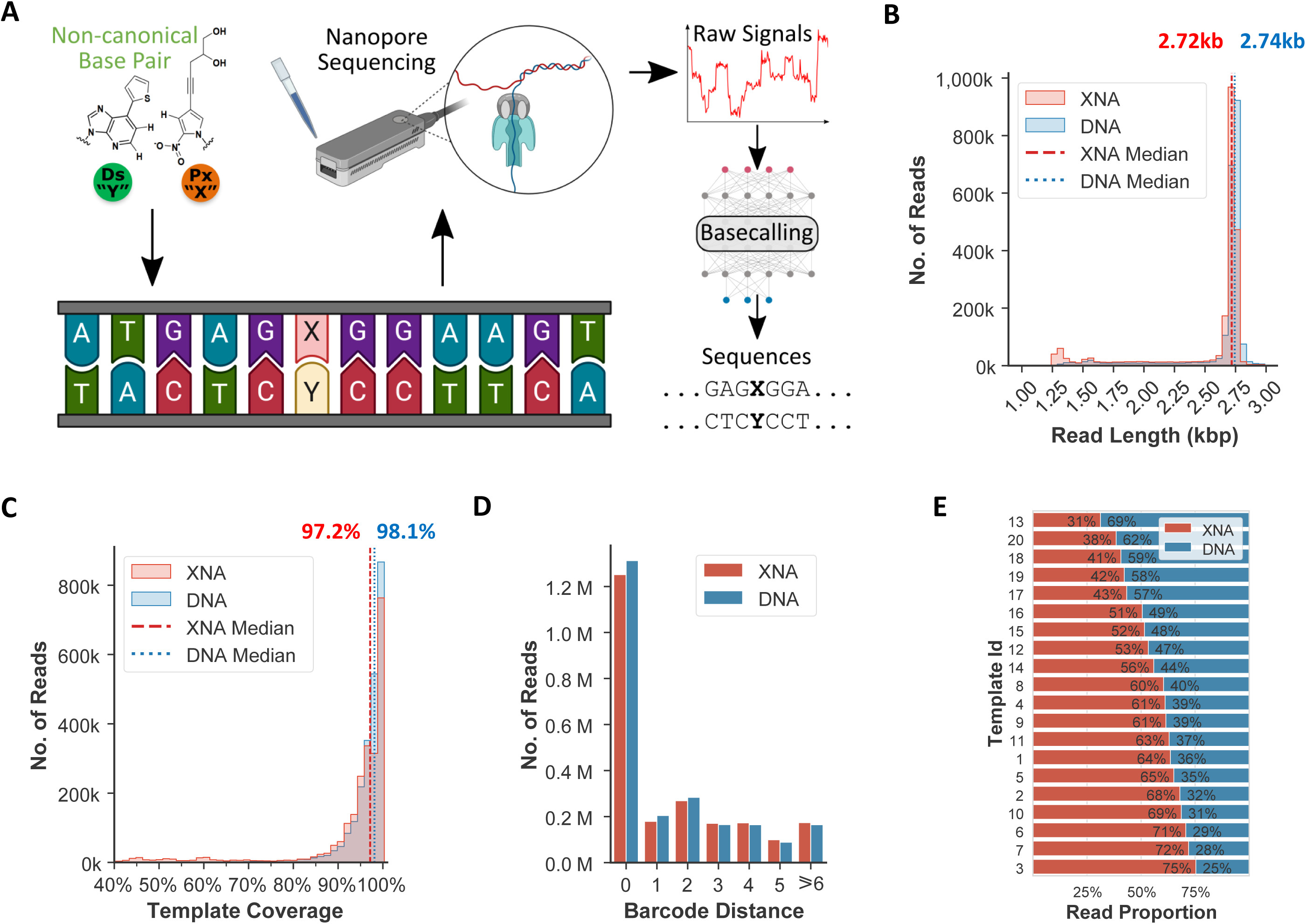
High throughput nanopore sequencing of XNAs. (A) Overview of workflow, showing the synthesis of XNA templates containing non-canonical basepairs (details in **Supplementary Figure 2**), nanopore sequencing, signal-level analysis and direct deconvolution of canonical and non-canonical bases. (B) Histogram of read lengths for XNA and control DNA libraries (dotted lines indicate median values). (C) Histogram of template coverage of template-specific region for XNA and control DNA libraries (dotted lines indicate median values). (D) Bar chart with read counts as a function of barcode distance (computed as edit distance of sequenced barcode to the closest reference barcode). (E) Read proportion between XNA and DNA sequences for each template, sorted by XNA proportion.

We then used a commercially available high-throughput nanopore sequencer (ONT MinION) to analyze the XNA molecules and control DNA (NCB replaced by canonical base), initially using standard sequencing protocols and basecalling pipelines (**Figure 1A**; **Methods**). The XNA sequencing process generated data continuously with no unexpected interruptions, with each flow cell yielding a total of >2 million raw reads (>5 Gbp of data), similar to yields from other sequencing runs with DNA templates^28^ (**Supplementary Figure 3**). However, XNA containing libraries had lower throughput as a function of time relative to the DNA libraries (mean=81%, 95% CI=69-99%), potentially due to a higher number of pore blockage and saturation events (39% vs 27% for control DNA). Despite this, in libraries containing equal concentrations of DNA and XNA templates, no strong bias was observed against XNA templates, with similar numbers of XNA and DNA reads being sequenced and successfully aligned to reference templates (2.3 vs 2.4 million reads). Read lengths for XNAs and DNAs were also very similar (median length 2.7kbp), with a substantial fraction of XNA reads (>75%) providing near full-length (±5%) template sequences (**Figure 1B**). A slightly higher fraction of XNA reads were shorter (1.2kbp) than the DNA reads recovered, due to incomplete fusion PCR^19^ in the XNA synthesis process (**Figure 1B**; **Methods**).

Alignment of reads to reference sequences provided similarly high coverage for XNA and DNA templates (median>97%; **Figure 1C**; **Methods**), though a higher fraction of XNA reads (33% vs 21%) exhibited incomplete coverage (<95%), potentially due to higher fragmentation rate or inability of conventional analytics to decode canonical bases correctly (particularly around NCBs). Demultiplexing successfully identified templates for a high proportion of both XNA and DNA reads (92.5% vs 93.1%, edit distance <5bp; **Figure 1D**), while the distribution of XNA and DNA reads also did not show a strong bias across the barcodes/templates where they were pooled together for sequencing (**Figure 1E**). In terms of basecalling accuracy, we noted that error rates were similar but slightly higher for canonical bases in XNA libraries versus control libraries (5% vs 3%; **Supplementary Figure 4**), consistent with the expectation that the presence of NCBs could induce basecalling errors in related signals. Taken together, these results highlight the ability to generate signal data for XNAs with high fidelity on a high-throughput nanopore sequencer, despite the system not being designed for it, and correspondingly the native basecaller’s inability to appropriately analyze NCB containing signals.

### Nanopore sequencing generates distinct electrical signals and reproducible errors near NCBs

After aligning reads to templates, we analyzed the corresponding raw electrical signal data measured by the nanopore sequencing device as a function of various sequence contexts (**Methods**). XNA signals showed NCB-specific divergence in contexts containing a single NCB (XNA01), as well as multiple NCBs that were close together (XNA13) or more spaced out (XNA16), and this pattern was seen for Ds as well as Px bases (upper and lower panels; **Figure 2A**). However, the largest difference was not necessarily observed when the NCB was in the middle of the kmer (**Figure 2A**), and specific patterns of divergence varied in different sequence contexts (**Supplementary Figure 5-7**). Overall, the average signal-level difference between XNA and DNA control sequences was observed to be greater near NCB positions (±3bp) than away from it (mean signal difference of 5.3 vs 0.7, median fold-change >6×), and the impact of sequence context was also seen in the form of larger variance for the distribution (43 in NCB context vs 0.4 in canonical context, **Figure 2B**). These results verify that the raw electrical signals generated from nanopore sequencing in the presence of NCB-containing XNAs can be notably distinct from those generated from an analogous DNA template.

**Figure 2.**
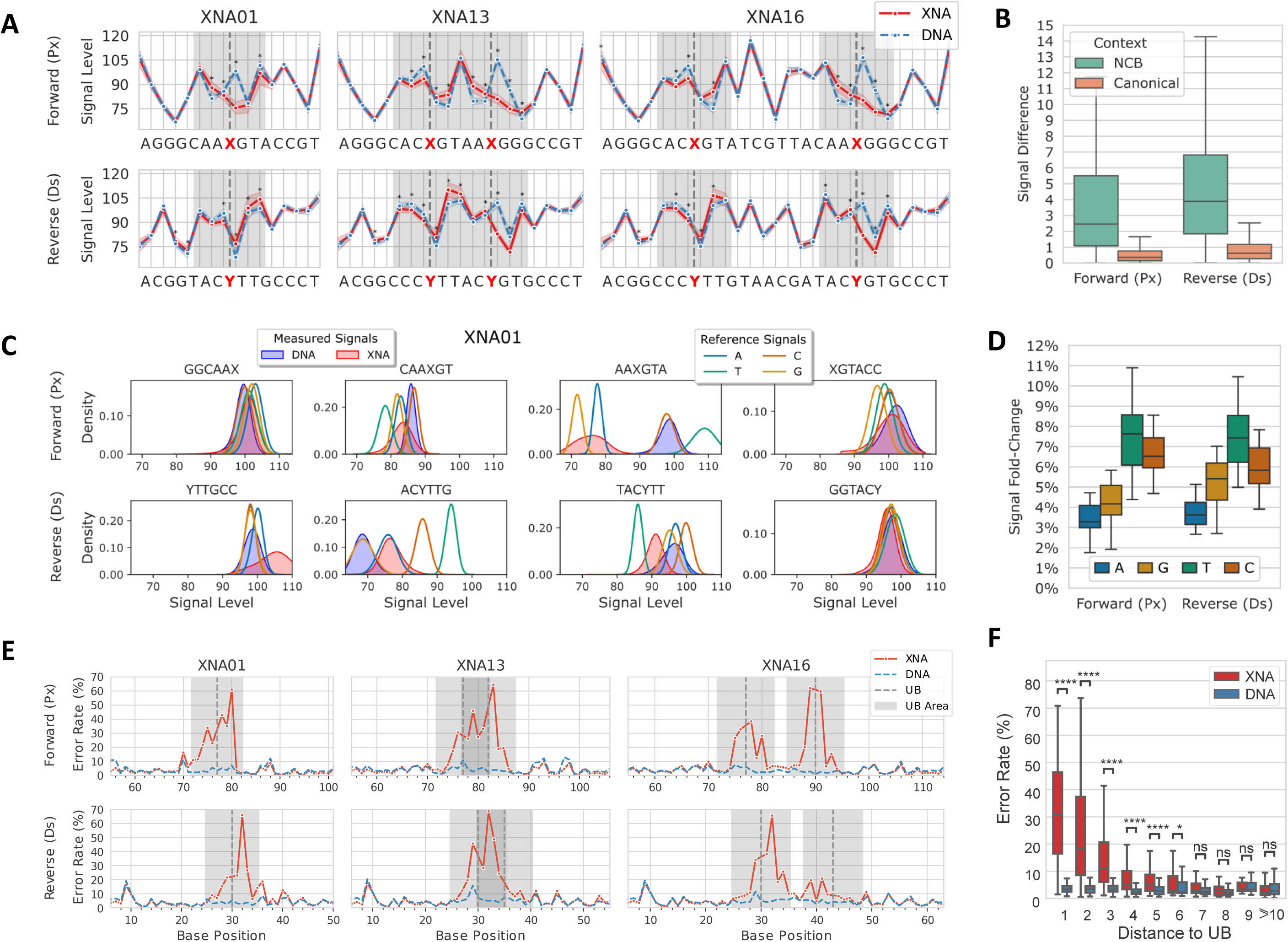
Nanopore sequencing generates distinct electrical signals and reproducible basecalling errors for non-canonical bases. (A) Average signal level comparison between XNA and control DNA reads for representative templates (other templates in **Supplementary Figure 5-7**). Vertical dashed lines indicate the location of the NCB, shaded areas indicate ±3bp windows around the NCB, and asterisks (*) indicate Bonferroni adjusted two-sided Wilcoxon p-value<0.05 and signal fold-change>2%. (B) Boxplots showing signal differences between XNA and DNA libraries for a template in the context of NCBs (±3bp window) and in canonical sequence contexts. (C) Signal level distributions based on read data for kmers containing NCBs and corresponding Nanopolish model distributions for kmers where the NCB is replaced by one of the canonical bases. (D) Boxplots showing average signal fold-change per template between read data for NCB kmers and corresponding Nanopolish model distributions for kmers where the NCB is replaced by one of the canonical bases. (E) Error rates per base position for selected templates with one or two NCBs. The NCB position is indicated by a vertical dashed line and shaded areas indicate 5bp windows around the NCB. (F) Boxplots showing error rates as a function of distance from the non-canonical base across all templates (“****”: One-sided Wilcoxon p-value<0.0001, “*”:p-value<0.05, “ns”: p-value>0.05). Summary statistics were computed based on sub-sampling of 5k reads per template and strand.

Subsequently, we investigated if signal-level distributions for NCB-containing kmers were distinguishable from reference distributions for kmers containing each of the canonical bases, based on a nanopore signal model^25^. In general, we noted a pattern where XNA-containing kmers exhibited signal distributions with distinct modes compared to canonical bases in several (but not all) kmers surrounding a NCB (similar signals when NCB is at end of the kmer and generally larger differences when NCB is at the center, **Figure 2C**). The fold-change per template between the non-canonical base and the mode for each canonical base had median value >5% for both Px and Ds bases (**Figure 2D**). These observations indicate that these NCBs are associated with electrical signal traces potentially distinguishable from all four canonical bases.

We next assessed the impact of signal-level differences on basecalling with the standard basecaller model and whether they introduced reproducible errors. In agreement with the signal-level comparisons (**Figure 2A**), XNA basecalling error rates also diverged from control DNA basecalling error rates in contexts containing a NCB (**Figure 2E**). Several templates exhibited very high error rates (>60%; **Figure 2F**) in the neighborhood of a single NCB (XNA01) as well as when multiple NCBs were present at varying distances to each other (XNA13 and XNA16). Notably, in contexts where canonical bases are between NCBs that are moderately spaced apart (>5 bp), we observed only a marginal effect for basecalling of those bases (XNA16, <10% error rate; **Figure 2E**), highlighting that the basecaller model can perform reasonably well even in regions flanked by NCBs.

Overall, in comparison to control sequences, the bases surrounding a NCB (±5 bp) showed substantially higher basecalling error rates (mean of 18% vs 3%; **Figure 2F**). The observed error rates are particularly high for the closest neighbors but quickly decrease with increasing distance away from the NCB (from 1 to 3 bp distance, median of 30%, 18% and 12%, respectively). In contrast, basecalling for positions far from the NCB was not impacted by its presence, obtaining error rates not significantly different from those for the control DNA templates (>6 bp; **Figure 2F**). Finally, for most templates (>90%) the native basecaller repeatedly miscalled the same canonical base more than 40% of the time (**Supplementary Figure 8**), emphasizing the consistency of the underlying signal data. These observations demonstrate that the performance of the standard basecaller is strongly and reproducibly impacted when deconvoluting NCB signals (i.e., bases affected by a NCB), while it is not impacted at positions sufficiently distant from NCBs, a feature that is ideal for bootstrapping the preparation of training data for learning a basecaller that can natively decode NCBs.

### Generation of a complex template library to enable training of XNA basecaller models

To obtain a new basecaller for XNAs (i.e. capable of natively basecalling canonical as well as non-canonical bases) we needed training data with enough reads and diversity of sequence contexts containing NCBs such that a neural network model with sufficient generalization capability could be developed. Since nanopore signal levels are known to be affected by the identity of at least the 6 adjacent bases in the pore during sequencing (**Figure 2A**), we designed a complex library where a NCB is flanked by 5-mers of the same sequence identity on both sides (4^5^=1,024 templates containing N_1_N_2_N_3_N_4_N_5_-M-N_1_N_2_N_3_N_4_N_5_ sequence, where: N_i_ = A, T, G, or C; M = Ds or Px; **Supplementary Figure 2**, **Supplementary File 2**-**3**), such that a rolling 6-nt window places the NCB in every position within the 6-mer resulting in all possible 6,144 single-NCB 6-mer sequence contexts being represented. Synthesizing such a large library would have been infeasible by employing the chemical synthesis approach that is typically used for obtaining high-fidelity XNAs, which was used for the initial proof-of-concept library here (20 templates). We therefore designed and developed a cost-effective enzymatic synthesis scheme to generate this much larger complex library (>50× in size; **Figure 3A**; **Methods**).

**Figure 3.**
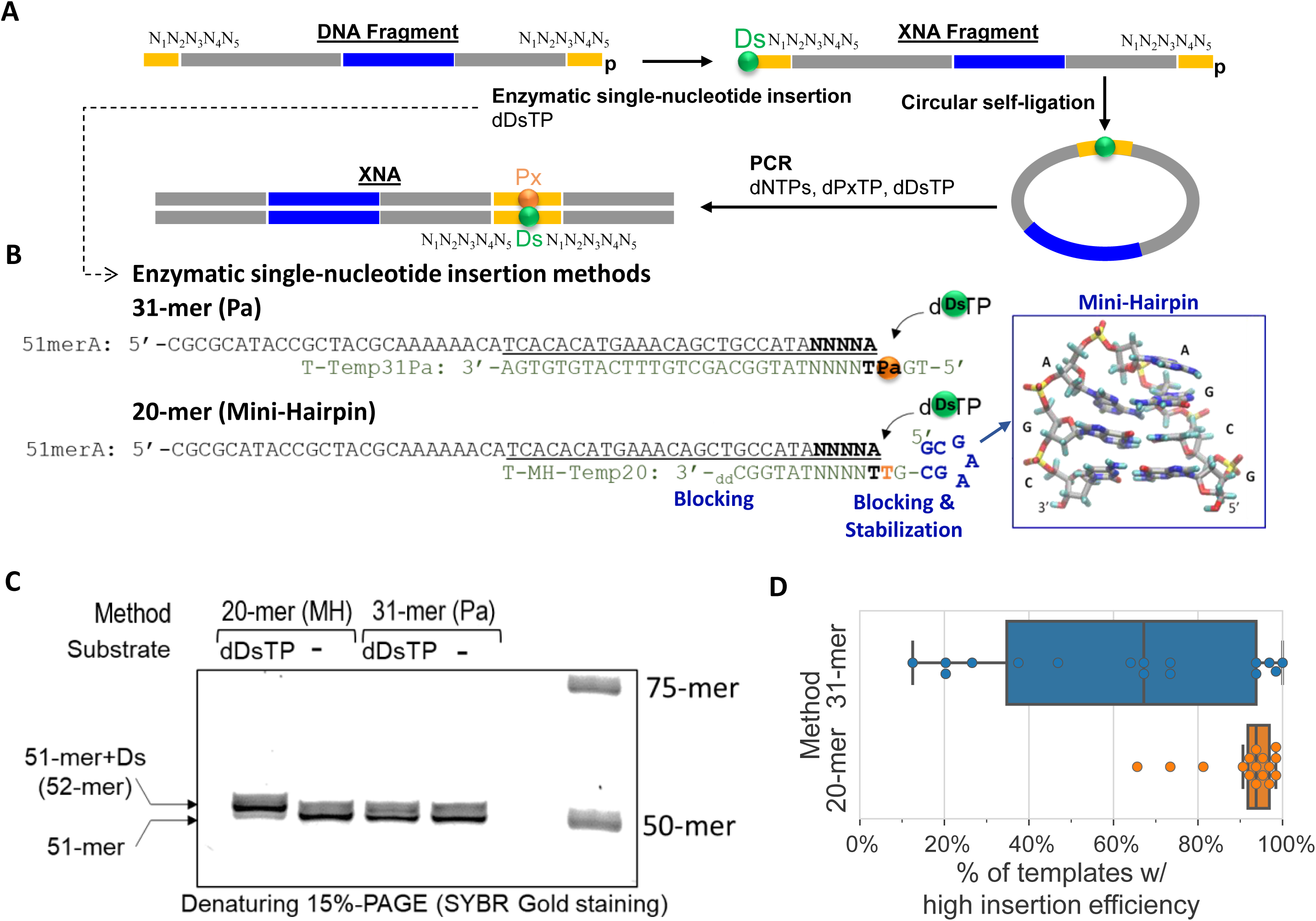
Generation of a complex XNA template library. (A) Schematic overview of XNA synthesis process, where DNA fragments phosphorylated at the 5’-end and containing the template and barcode sequences undergo enzymatic single-nucleotide insertion of Ds at 3’-terminus. The resulting XNA fragments are then subjected to circular self-ligation followed by PCR using canonical and non-canonical base substrates to obtain the double-stranded XNA. (B) Figure depicting the two alternative strategies that were explored for enzymatic single-nucleotide insertion, using a 31-mer (Pa) and a 20-mer (mini-hairpin) template. (C) Gel electrophoresis results for single-nucleotide insertion products using the two alternative strategies. (D) Boxplots depicting Ds insertion efficiency per pool of templates for the two alternative strategies. Insertion efficiencies were estimated via replacement PCR and Ion PGM sequencing.

The proposed scheme consists of enzymatically inserting the Ds base at the 3’- end of a fixed DNA sequence. For this insertion, we explored two methods using different templates (31-mer and 20-mer; **Figure 3B**). In the first method, we used 31-mer templates containing the Pa nucleotide (more stable alternative to Px). However, only a moderate yield for the Ds-inserted product could be obtained from the reaction mixture (even after purification by denaturing gel electrophoresis; T-Temp31Pa; **Figure 3C**). Subsequently, we designed a second method, using a 20-mer template with a mini-hairpin DNA sequence to stabilize the structure at the insertion target site (20-mer; **Figure 3B**). This method resulted in higher Ds insertion efficiency (T-MH-Temp20; **Figure 3C**).

To assess the insertion efficiency of the proposed enzymatic synthesis scheme, we estimated the purity of Ds insertion per pool of templates (**Figure 3D**; **Methods**), where each pool was comprised of subsets of templates with the same first and fifth bases in the target sequence (**N_1_***N_2_N_3_N_4_***N_5_**, 16=4**^2^** pools with 64=*4^3^*fragments each; **Supplementary Figure 9**). Considering the proportion of pools with high NCB retention rate (>85%), the 20-mer (mini-hairpin) method provided a substantial improvement over the 31-mer (Pa) method (median of 94% *vs* 67%; **Figure 3D**). Overall, a great majority of XNA templates (91%) synthesized via the mini-hairpin method were estimated to have high-purity (>85%). These results emphasize how an enzymatic synthesis scheme with the right design can still enable generation of a substantially more complex template library reliably, and with purity high enough that it could enable effective training of a XNA basecaller model.

### Bootstrapping and data augmentation enables training of an accurate generalized XNA basecaller

The complex template library was sequenced on a MinION sequencer, generating >6 million reads, with similar level of success as the proof-of-concept library in terms of long read lengths, near full-length template coverage and successful barcode identification (**Supplementary Figure 10**). The distribution of reads across templates was found to be relatively similar, with hundreds of reads being available per template for most templates, after preprocessing and filtering of short reads (**Supplementary Figure 11**; **Methods**). XNA training data was prepared by pre-processing reads selected for training by splitting them into signal chunks and attributing ground-truth sequence based on the library reference template sequence, discarding chunks that matched poorly to the reference (**Supplementary Figure 12**; **Methods**). A significant fraction of reads was discarded in this way in an initial round of training (29%), as the baseline DNA model is prone to high error rates within NCB regions (**Figure 2E-F**, **Supplementary Table 1**). Nevertheless, the resulting model was able to substantially improve basecalling in bases adjacent to the NCB (93.8%, Round 1) versus the baseline model (75.6%, Round 0; **Figure 4A**). The updated basecaller model was used to repeat the pre-processing of reads, resulting in much fewer reads being discarded (11%), and retraining of a basecaller model with improved accuracy for calling NCBs and neighboring bases (>94% in Round 2; **Figure 4A**). Further rounds of train data preparation and basecaller fine-tuning had limited impact on the number of reads added (<1%) and improvement of model performance (Round 3, **Figure 4A**), indicating that one round of bootstrapping was sufficient to obtain an accurate model for the complex library.

**Figure 4.**
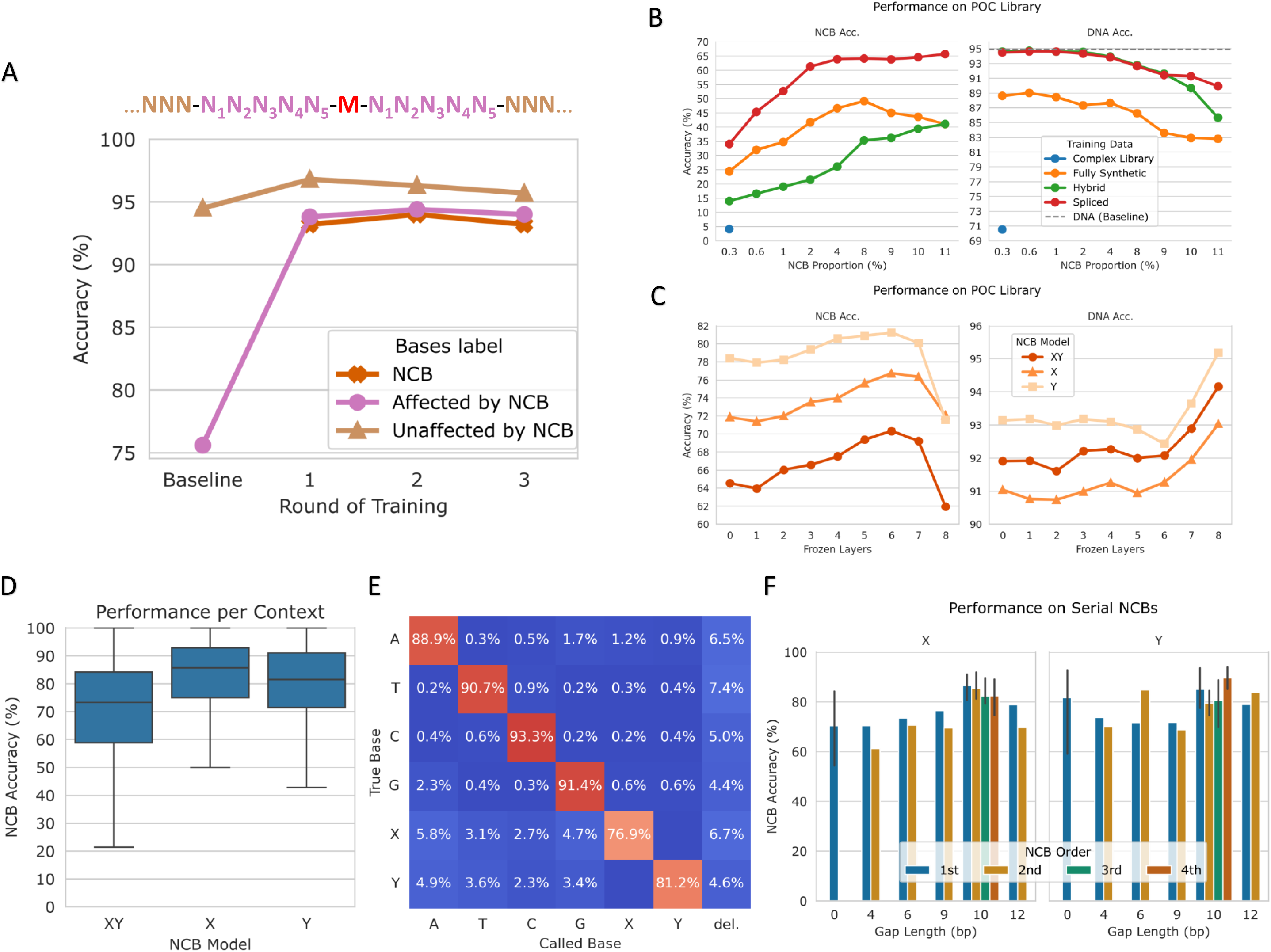
Training of a high-accuracy XNA basecalling model. (A) Line plot depicting base-level accuracy per round of training for non-canonical bases, canonical bases within 5bp of a non-canonical base and more distant canonical bases. The baseline round of training depicts results using the Bonito Super Accuracy DNA model. (B) Line plots depicting performance on the proof-of-concept library using basecaller models trained with different sets of real and artificial reads, with increasing proportion of NCBs per read. The following figures are based on models trained with spliced data and 9% NCB proportion. (C) Line plots depicting performance of models trained jointly for both NCBs or separately, with increasing number of layers frozen. Subsequent analyses are based on models with 6 frozen layers. (D) Boxplots depicting performance of the final model, based on the complex library for testing with diverse templates and held-out reads. (E) Basecalling confusion matrix for NCBs and canonical bases. The last column represents deletion errors. (F) Barplots (error bars represent 100% percentile interval) depicting performance analysis for reads with multiple NCBs. NCB proximity, number and order did not seem to have a strong impact on NCB accuracy, indicating that the final trained basecaller is robust in handling reads with multiple NCBs.

Model training based directly on XNA reads from the complex library is expected to have low generalizability as the sequence context surrounding the NCB containing kmers is fixed. Correspondingly, testing on reads from the proof-of-concept 20-member library demonstrated low accuracy for both NCBs and canonical bases (5% NCB accuracy, 71% DNA accuracy; Complex Library, **Figure 4B**). To overcome this limitation, we supplemented model training with different data augmentation approaches, allowing us to generate artificial reads with raw signals corresponding to NCB containing kmers placed in diverse sequence contexts. Initially, fully synthetic signals were simulated based on a signal-level model trained on real signal data from the complex library (Fully Synthetic, **Supplementary Figure 13**; **Methods**). This approach notably improved NCB and DNA accuracy (24% and 89% respectively) relative to the model trained directly on the complex library, where the proportion of NCBs was kept constant (0.3%; Fully Synthetic, **Figure 4B**). However, as DNA accuracy was lower than the baseline DNA model (94%), we investigated a hybrid approach containing a mixture of real DNA signals with simulated XNA signals that were generated by replacing specific positions in real DNA chunks with synthetic signals representing NCB kmers (Hybrid, **Supplementary Figure 13**; **Methods**). This approach provided slightly lower NCB accuracy relative to the fully synthetic version, but higher DNA accuracy that matched the baseline model (14% and 94% respectively; Hybrid, **Figure 4B**). The final strategy that was tested involved splicing real DNA and XNA signals together to generate artificial XNA reads with high context diversity while retaining the fidelity of real sequencing data (Spliced, **Supplementary Figure 13-14**; **Methods**). Training based on these spliced chunks resulted in substantial improvement in NCB accuracy without compromising DNA accuracy (34% and 94% respectively), indicating that it provides the most robust and generalizable approach for data augmentation and model training (Spliced, **Figure 4B**).

Leveraging the flexibility of this approach for data augmentation, we next explored various ways to improve model performance. By increasing the proportion of NCBs in each generated read, in contrast to using a single NCB per read, substantial gains in NCB accuracy were noted (∼2× to 64% at 4% NCB) with only a slight reduction in DNA accuracy (∼1%; **Figure 4B**). While NCB accuracy can be further improved, this comes with diminishing returns at higher NCB proportions (66% NCB accuracy and 90% DNA accuracy; trained with 11% NCB proportion, **Figure 4B**). As baseline models are well-adapted to call DNA bases, we next explored a *domain adaptation* approach by preserving these pre-trained weights for some layers during fine-tuning (**Methods**). Successively freezing layers of the basecaller architecture up to layer 6 (out of 9; **Supplementary Figure 15**) helped boost model accuracy without substantially altering DNA accuracy (70% NCB accuracy for XY model with 6 frozen layers; **Figure 4C**). Not surprisingly, models trained separately for each of the NCBs achieved even higher accuracy (77% for X and 81% for Y), which could be leveraged for highly accurate joint basecalling when both strands of the XNA are sequenced in duplex mode (**Figure 4C**).

Using the complex library to evaluate model performance across diverse sequence contexts further confirmed that high NCB accuracies are typically obtained by our model across templates (median accuracy=86% and 82% for X and Y, and 73% for XY model; **Figure 4D**). In addition, this comes with very high specificity (99%), and thus canonical bases are rarely miscalled as one of the NCBs (**Figure 4E**). Instead, a significant proportion of the errors involve NCBs being missed (i.e. a base deletion), similar to what is seen in nanopore basecalling models for canonical bases as well (4-7%; **Figure 4E**). In addition, NCBs are more likely to be miscalled to a canonical base, relative to the canonical-base-to-canonical-base miscall rate (2-6% vs 0-2%; **Figure 4E**), suggesting that our models are conservative. Nevertheless, our models achieve good recall rates (77% and 81% for X and Y models, and 75% for XY model). Finally, we did not observe a strong influence from the order of NCBs in a template, or the number of canonical bases between consecutive NCBs, in terms of X and Y basecalling accuracy (**Figure 4F**). Moreover, model performance was also comparable for templates containing multiple NCBs (including a 6-mer with two NCBs), despite these configurations not being present in the training data (**Figure 4F**), highlighting the generalization capability of the model. Overall, these results highlight the ability to learn a high-accuracy generalized direct basecalling model for XNA sequences using bootstrapping and data augmentation to increase data complexity.

## Discussion

In this work, we provide the first demonstration of direct high-throughput sequencing and deconvolution of non-canonical bases that can be readily performed on a commercial sequencer. This effectively expands the set of bases that can be natively read by nanopore sequencers paving the way for in-depth exploration of other classes of non-canonical bases as well^29,30^. The workflow used here can serve as a general template for this, including (i) systematic assessment of the ability of a nanopore sequencing system to process NCB containing templates with high fidelity (section 1), (ii) evaluation of the resulting signals for distinctness and reproducibility (section 2), (iii) synthesis of a complex training library using a mini-hairpin sequence to enable high efficiency (section 3), and (iv) training of a high-accuracy model using bootstrapping and signal splicing to boost data complexity (section 4). More specifically, our results highlight that ONT’s nanopore sequencing device can be applied out-of-the-box to XNAs containing multiple Ds-Px unnatural base pairs, processing them to generate raw electrical signals without significant disruption, and with only a slight reduction in throughput to produce >2 million reads and >5Gbp of data in a single MinION run (**Supplementary Figure 3**). The resulting read sets were frequently near full-length (>75% within 5% of expected length) and showed no notable biases in data generation efficiency compared to their DNA controls (**Figure 1B-E**). These results bode well for the use of nanopore architectures as versatile systems for natively reading and decoding diverse non-canonical bases.

The raw signal data, however, does not directly provide nucleotide base identities. To assess if it could be used for training a basecaller that could directly deconvolve signals into bases, we first systematically assessed the distinguishability, fidelity and reproducibility of the signal data. We found that the native basecaller was robust enough to produce DNA reads that were largely accurate, except in regions surrounding the NCB (**Figure 2F**). This enables accurate alignment of signal data to reference bases, and consequently preparation of training data for machine learning with NCB containing XNA signals. Notably, signal data around a NCB (±3 bp) is substantially different relatively to control DNA reads (median fold-change >6×; **Figure 2A-B**), and these differences are not apparent outside the NCB region. We further noted that the native basecaller makes reproducible errors in XNA containing reads indicating that the underlying signal patterns could be reproducibly re-interpreted with an appropriately trained basecaller (**Supplementary Figure 8**).

Training a basecaller that directly deconvolves NCBs along with canonical bases requires training data that contains signals from XNA reads with known sequences in a sufficiently complex library. While chemical synthesis approaches have high efficiency, they can be prohibitively expensive for generating large-scale libraries. Enzymatic synthesis approaches are generally known to provide modest insertion efficiencies (7%-73%)^30^ at lower cost. Here we demonstrated that a complex XNA library (n=1,024 templates) can be generated with high insertion efficiency (>85%) by designing a custom insertion template and a mini-hairpin to stabilize the NCB at the target site (**Figure 3C-D**). A similar strategy could be useful for designing complex libraries for other NCBs as well. We chose to design our library to capture signals generated by all 6,144 single-NCB 6-mers sequence contexts (**Figure 3A, Supplementary Figure 2**). This improves on previous work based on 4-mers^30^, and is expected to benefit model training by capturing greater sequence context diversity in critical bases impacting signal data^31^. In principle, generating data with 6-mers containing multiple and consecutive NCBs would also benefit model training. However, this is currently infeasible with even modest levels of efficiency^7^ and therefore a limitation of our work is that we are currently unable to assess the capabilities of our model with such XNAs. Since directly sequencing XNAs containing consecutive NCBs would expand the utility of this model for synthetic biology applications, such as while creating XNA enzymes or genomes, we aspire to address such limitations when advances in the synthesis process allows the generation of data with such characteristics. Noteworthily, prior work using different classes of NCBs and a non-commercial nanopore demonstrated that having homopolymers with NCBs increases premature nanopore dissociation, resulting in many incomplete reads^32^. Thus, in order to successfully sequence NCB homopolymers, future adaptations to nanopore chemistry or the NCB structure itself might be required.

Despite the sequence diversity in the immediate neighborhood of the NCB in our complex library, the XNAs that we sequence still have a predictable sequence backbone (**Supplementary Figure 2**). Not surprisingly therefore, using this read data directly provides a high-accuracy model for reads from the library (94% for UBs; **Figure 4A**), but does not generalize to other sequence contexts. Nevertheless, this initial model based on bootstrapped learning allows us to align signal data from the complex library to its corresponding reference bases accurately and thus enables further robust training. As generalizability is a common problem for training XNA models with limited training data complexity, we explored several strategies for data augmentation including generating fully synthetic signals, as well as signals that splice combinations of real and synthetic data. Our results suggest that retaining signals from real XNA reads while embedding them in diverse DNA read contexts provides higher accuracy relative to synthetic data (>80%) and robust generalizability for the model (**Figure 4B-F**). The models reported here also exhibit very high-specificity (99%) relative to prior work^30^ (80-93%), while demonstrating the capability for the first time to directly deconvolve signals and read NCBs and canonical bases in an XNA. Our strategy for data augmentation is versatile and can be adjusted to different requirements. For example, lower NCB proportions can be used when prioritizing DNA accuracy over NCB accuracy, while NCBs that were not synthesized together can be incorporated into the same artificial read, to improve model accuracy for more densely packed NCBs and different classes of NCBs. Notably, data augmentation might be even more important as we seek to expand the alphabet of bases that can be directly read by nanopores ^5,30^, and the corresponding demands for library complexity go up rapidly.

With ongoing advances in nanopore sequencing technology, there are several opportunities to further enhance capabilities to sequence NCB-containing templates by directly optimizing the design of motor enzymes, pores and signal modalities (e.g. variable-voltage^33^). Additional measurements (e.g., duplex-basecalling, translocation time and backwards step) could also substantially aid in closing the gap between NCB and canonical basecalling accuracy. Since the signal trace corresponding to a particular nucleobase sequence greatly influences basecalling performance, a deeper understanding of the underlying physicochemistry could help to significantly improve basecalling of XNAs. This knowledge would enable the development of NCBs that produce electrical signals more distinct from those produced by canonical bases, thereby reducing or eliminating ambiguities that often cause sequencing errors. Advances in the deep learning architecture of basecallers can also be beneficial for modeling NCB signals, especially if future models become more generalizable without needing extensive amounts of training data.

High-throughput NCB sequencing can, in turn, pave the way for advances in various fields. For example, DNA storage applications can greatly benefit from the much higher information density that an expanded alphabet enables^17,29^. In synthetic biology, the development of new UBs, new XNA synthesis methods, and the creation of synthetic genomes that exploit UBs can be transformed by the ability to directly sequence XNAs. Finally, the development of NCB-containing aptamers^21,34^ and other DNA therapeutics would benefit from easier access to direct XNA sequencing. More broadly, we hope that the proof-of-concept shown in this work further accelerates the use of nanopore sequencing techniques to interrogate the role of non-canonical basepairs across various domains of life, particularly for viral genomics^11^ as well as for sequencing tRNAs^9^.

## Methods

### Preparation of XNA fragments containing canonical and non-canonical bases

XNA fragments containing Ds or Diol-Pa were chemically synthesized with an automated DNA synthesizer (H-8 SE DNA/RNA synthesizer, K&A Laborgeraete) using Ds and Diol-Pa phosphoramidite prepared in-house^6,35^ and commercially available canonical base phosphoramidites (Glen Research), or purchased from Gene Design. In-house synthesized oligonucleotides were purified by denaturing gel electrophoresis, after deprotection with concentrated ammonia solution. The non-canonical base substrates, dDsTP, dPxTP, and dPa’TP, were prepared in-house as described previously^6,36^. DNA fragments consisting of canonical base sequences were purchased from Integrated DNA Technologies (IDT) and used to prepare short and long XNAs (**Supplementary File 4**).

### Preparation of proof-of-concept 20-member XNA Library

Using chemically synthesized DNA fragment sets, we first prepared short XNAs with barcode sequences, through primer extension in the presence of dPxTP and then PCR amplification in the presence of dDsTP and dPxTP as additional non-canonical base substrates, as described previously^19^. Combinations of Ds-containing templates and the barcode fragments are summarized in **Supplementary File 5**. The presence of non-canonical bases at specific positions in the PCR-amplified short XNAs (∼97% purity) was confirmed by an in-house Sanger gap sequencing method^19^. For nanopore sequencing analysis, we prepared long XNAs by fusion PCR as described previously^19^, in the presence of dDsTP and dPxTP, using three DNA blocks: short XNAs prepared as described above, left-arm DNA fragments, and right-arm DNA fragments. The presence of non-canonical bases at specific positions was confirmed by an in-house Sanger gap sequencing method^19^. The control long DNAs, where the non-canonical bases were replaced by canonical bases, were prepared by fusion PCR in the absence of non-canonical substrates, using the short DNA blocks with chemically-synthesized templates comprising of canonical base only (for XNA01–XNA16) or prepared by replacement PCR in the presence of dPa’TP^22^ (for XNA17–XNA20).

### Preparation of complex XNA library

#### Ds insertion

Synthetic 90-mer oligonucleotide pools (oPools^TM^ Oligo Pool, **Supplementary File 2-3, Supplementary Figure 9**) were purchased from IDT, as mixtures of 50pmol fragments. DNA fragments with phosphorylation at 5’-end (ordered with phosphorylation or phosphorylated by T4 DNA polynucleotide kinase followed by denaturing-gel purification) were directly used for single-Ds nucleotide insertion by exonuclease deficient Klenow Fragment DNA polymerase (KFexo-, New England Biolabs).

In the initial trial, each 90-mer DNA pool was annealed with the corresponding 31-mer template (5’-TG(Diol-Pa)NNNNNTATGGCAGCTGTTTCATGTGTGA-3’, N=A, G, C, or T), positioning Pa opposite the site intended for Ds insertion. The Ds insertion was performed through the incubation of the 90-mer DNA pool (2μM) with 31-mer template (20μM) in the presence of 100μM dDsTP and 0.2U/μl KFexo-in the reaction buffer (25mM Tris-HCl pH 7.5, 10mM MgCl_2_, 1mM DTT, 50mM NaCl) for 60 min at room temperature. The inserted products were purified by denaturing PAGE. However, we found that Ds insertion rate was relatively low, so we designed a new type of template for Ds insertion using a 20-mer containing a mini-hairpin DNA sequence at 5’ terminus, X-MH-temp20 (5’-<u>GCGAAGC</u>GTNNNNNTATGG(ddC)-3’, N=A, G, C, or T, underlined: mini-hairpin sequence). The design was based on three points: (1) the mini-hairpin sequence at the 5’-terminus allowed specific and stabilized hybridization to the 3’-region of 90-mer pool, (2) the use of 2’,3’-dideoxy C at the 3’-terminus prevents additional Ds insertion, and (3) Ds insertion opposite T is allowed since we add only dDsTP (**Figure 3A-B**). As expected, the Ds insertion in the presence of the 20-mer template was much improved compared with that in the presence of the 31-mer (**Figure 3C-D**). Finally, the Ds insertion to 90-mer pool was performed through the incubation of the 90-mer DNA pool with 20-mer template (75μM) in the presence of 100μM dDsTP and 0.5U/μl KFexo-in the reaction buffer for 30 min at 37°C.

#### Circligation

The prepared Ds-inserted DNA pools (around 1.5 to 2.5 pmol) were subjected to self-ligation (10μl) by 50 Units of Circligase II (Lucigen) in the circligation buffer supplemented with 2.5mM MnCl_2_ and 1M Betain for 16 h at 60°C, followed by heat inactivation of the ligase at 80°C for 10 min. The unligated fragments were removed by treatment with Exonuclease I (NEB) and Exonuclease III (NEB) at 37°C for 45 min, followed by heat inactivation of the nucleases at 80°C for 20 min.

#### Preparation of short and long XNA pools

The reaction aliquots were subjected to 12-cycle PCR amplification in the presence of dDsTP and dPxTP, using two primers, pUC19rev-47 and pUC19fwd-42. The PCR-amplified products (short XNA pools) corresponding to 135-bp (without Ds-Px) /136-bp (with Ds-Px) were purified by denaturing PAGE, followed by fusion PCR to prepare long XNAs, as described above. The presence of non-canonical bases at specific positions in the PCR-amplified short/long XNA pools were confirmed by an in-house Sanger sequencing method supplemented with ddPa’TP^19^ or Ion PGM deep sequencing after replacing the non-canonical bases with canonical bases^22^.

### Nanopore library preparation and sequencing

XNA library samples were sequenced on Oxford Nanopore Technologies (ONT) MinION Mk1B or Mk1C devices using MinION R9.4.1 flowcells. These samples were prepared using the ONT Ligation Sequencing Kit (SQK-LSK109 or SQK-LSK110) following standard library preparation protocol instructions provided by the vendor. The nucleic acid concentration of each sample was measured using the Qubit^TM^ dsDNA HS kit on a Qubit^TM^ 3 Fluorometer (Invitrogen).

In total, we ran five different synthesis and sequencing batches: 1) XNA+DNA #01-16; 2) XNA #17-20; 3) DNA #17-20; 4) Complex library; 5) Complex library subset (**Supplementary Figure 2**, **Supplementary Figure 9**). Run #5 was needed to replenish some of the templates (n=214) with low read counts in run #4.

### Nanopore data pre-processing

To prepare XNA sequence reads for raw nanopore signal analysis and basecaller training, we used Guppy (v4.0.14) with the dna_r9.4.1_450bps_hac profile for initial basecalling. The output basecalled sequences were then aligned to template references using Minimap2^37^ (v2.17-r941; parameters -x map-ont -c -secondary=no -w 5). For alignment we used only template-specific regions as references, i.e. the sequences corresponding to the left and right arms were omitted from the reference targets, as this helped to avoid misalignment to the wrong target. After alignment, the barcode region was located and used to compute the number of mismatches with the reference barcode to obtain edit distances. Reads with low template coverage (<85%), high barcode distance (>5) or unexpected read length (<1K or >3k) were filtered out, before counting the final number of reads per template (**Supplementary Figure 11**). Nanopolish^25^ (v0.13.0; parameters eventalign -samples -scale-events -print-read-names) was used to obtain the mapping between read bases and raw signals.

### XNA training data preparation

We randomly sampled 316k reads from the complex XNA library sequencing batches, of which 297k reads were used for training and 19k reads for validation, with respective medians of 170 and 10 reads per template and strand. We prepared the input for training the basecaller by first dividing the raw signals into chunks with a chunk size of 3600 and an overlap of 900 signal points. Each chunk was then basecalled and mapped to the library reference to retrieve the ground-truth sequence. We retained only the chunks that contained a NCB and had at least 90% coverage and accuracy relative to the aligned reference (**Supplementary Table 1**). Each chunk of signals 𝑆 = {𝑆_1_, …, 𝑆_𝑐ℎ𝑢𝑛𝑘𝑠𝑖𝑧𝑒_} was normalized via SMAD (Scaled Median Absolute Deviation) normalization^38^, as defined below:

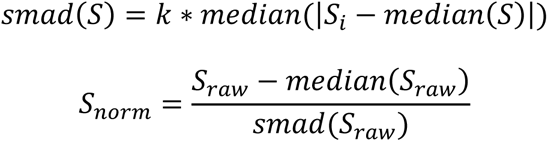

where, for consistency with the Gaussian distribution, we used 1.4826 as the scale factor 𝑘. We employed SMAD normalization to comply with Bonito’s signal pre-processing step before basecalling using its pre-trained models.

### XNA basecaller architecture and training

We used Bonito^39^ (v0.5.0) Super Accuracy (SUP) as the backbone architecture for our XNA basecaller models. We chose Bonito for the backbone because it is the state-of-the-art in terms of basecalling accuracy^40^, and ONT’s recommended basecaller for training new models and method development. We made certain adaptations to Bonito’s implementation and basecaller architecture (**Supplementary Figure 15**) to enable training with data containing a six-letter alphabet. The baseline model, trained solely on DNA data, was used as the default set of initial weights for fine-tuning with the XNA data. After round 1, rounds 2 and 3 of the bootstrapping strategy used the weights from the previous round to extract training chunks and initialize the model for fine-tuning (**Figure 4A**). Unless otherwise specified, we fine-tuned the weights from all layers. However, in some cases, we partially froze the network and only fine-tuned the weights from the top layers (**Figure 4C**). We fine-tuned the network for 5 epochs with a learning rate of 5e-4, applying dropout rates of 50% to the top layer and 5% to any remaining unfrozen layers.

### Performance evaluation

To comprehensively evaluate our basecaller models, we assessed their performance on both XNA libraries (proof-of-concept and complex). For performance evaluation over the proof-of-concept XNA library we randomly sampled 10k reads, consisting of 250 reads per NCB and template (**Supplementary Figure 16**). For the complex library we randomly sampled 40k reads (median of 20 reads per NCB and template). Each model was used to basecall the two test sets, and evaluation was conducted on the reads successfully aligned to reference templates. To evaluate how well our models can basecall canonical bases (ACGT) and NCBs (XY), we calculated base-level accuracy and reported the average accuracy per template and strand for each group of bases (canonical and non-canonical). We categorized canonical bases within 5 bases of a NCB into a separate group (i.e. NCB-adjacent bases) due to their susceptibility to the influence of the NCB at the signal level and, consequently, in terms of basecalling. We computed recall and specificity metrics considering NCBs as positives and canonical bases as negatives.

### Refined signal-to-sequence alignment

We used Nanopolish on the reads from the complex XNA library to obtain an initial signal-to-sequence alignment, which was subsequently refined. In order to obtain an alignment with higher quality, we used the sequences basecalled by the XNA model from round 2 of the bootstrapped training procedure and we substituted the NCBs in the reference sequences with the canonical base that minimized the average standard deviation per kmer (A for both X and Y). We filtered out reads with alignments that have coverage <75%, a position with signal count >50, an average variance per position >20 or a signal match score >2. We also computed an alternative score by matching the signals to a reference sequence without the NCB, representing the mis-insertion case, and discarded the reads where the difference between the alternative and original scores was less than −0.25, capping discarded reads at 15% of the total. The signal match score for a given sequence of signals grouped by kmer is the average z-score based on kmer model mean and standard-deviation and is defined as:

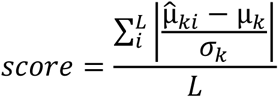

where L is the sequence length, 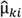 is the observed mean signal level for kmer *k* at position *i* of the sequence, µ_𝑘_ is the model mean signal level for kmer *k* and 𝜎_𝑘_ the model standard deviation. We used the kmer signal model parameters employed by Nanopolish.

Since Nanopolish was designed solely for canonical bases, the output alignment is not very accurate around non-canonical bases, necessitating additional steps to improve it. After filtering the reads, we employed dynamic time warping (DTW)^41^ in two iterations to refine the signal-to-sequence alignment in the region comprising the NCB kmers and the 5 kmers before and after. Initially, we substituted both the NCBs with the canonical base “A” (since it minimized kmer standard deviation for both X and Y) and used the model mean signal level as the reference signal for each kmer. Subsequently, we updated the reference corresponding to the NCB kmers with the observed mean signal level, computed from the signals aggregated across all reads after the first iteration. We ran the DTW algorithm with Euclidean distance and an asymmetric step pattern, which ensures that every signal is always matched to a single kmer. Additionally, to enforce that each kmer would be matched to at least three signals, we repeated each reference signal three times.

### Synthetic signal generation

We simulated nanopore signals using the modeled signal distribution (mean and standard deviation) of each 6-mer. For DNA kmers, we used the model available with the Nanopolish tool. However, for XNA kmers, we had to generate our own model covering the NCBs explored in this work (Px-Ds). We modeled the XNA kmers by first applying our refined signal-to-sequence alignment on the sequenced reads from the complex XNA library. We then grouped the observed XNA signals per 6-mer, removed outliers (±3 × 𝑠𝑡𝑑𝑒𝑣), and extracted the mean and standard deviation from the remaining samples (**Supplementary File 6**).

Using the 6-mer signal model for DNA and XNA, we simulated the signals for a specific kmer by sampling from a truncated normal distribution, with the kmer model mean and standard deviation as parameters, and then adding Gaussian noise^42^. The generation of a synthetic signal 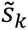 for kmer *k* is defined as follows:

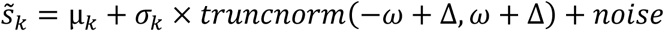

where 𝜔 and Δ are the window size and shift value, respectively, for defining the truncation lower and upper bounds. We used a window size of 1.5 and a shift value randomly chosen from −0.5, 0 and 0.5. The Gaussian noise was also sampled from a truncated normal distribution, with a standard deviation scale randomly selected within the range of 0 to 1 and lower and upper bounds of ±3. The random parameters (window shift and noise standard deviation) vary between reads but are chosen once and fixed for each read.

### Fully synthetic XNA chunks

We used the DNA training data from Bonito as the source for the reference sequences for which we generated fully synthetic nanopore reads. Adhering to a target NCB proportion for the new XNA reads, we randomly selected multiple positions from the source reference sequence to be replaced by X or Y, ensuring that no 6-mer contained more than one NCB by maintaining a minimum of 5 bases between selections. We also used Bonito’s data to determine the number of signal points each kmer instance should have. To estimate this information, we segmented the chunk of raw signals corresponding to the source sequence into kmers. Finally, we normalized the resulting chunk of signals using SMAD normalization.

### Hybrid XNA chunks

We created hybrid reads by replacing specific signals in DNA chunks from Bonito training data with synthetically generated XNA signals. Given a DNA chunk and its reference sequence, we selected multiple positions from the sequence to be replaced by NCBs and then substituted the signals corresponding to these positions with nanopore simulations for the NCB kmers resulting from the replacement. To determine which signals correspond to the replaced positions, we segmented the DNA signal chunks into kmer instances employing DTW with the model mean signal level serving as the reference signal for each kmer. Similar to the final step of our refined signal-to-sequence alignment, we ran the DTW algorithm with Euclidean distance, an asymmetric step pattern, and repeating each reference signal three times.

### Spliced XNA chunks

To achieve more diverse sequence contexts while maintaining high nanopore signal fidelity, we assembled spliced reads by replacing specific signals in DNA chunks from Bonito training data with XNA signal slices extracted from our nanopore XNA reads. We used the chunks from the complex XNA library, obtained with the basecaller model from round 2 of bootstrapped training, as the source for the signal slices for NCB kmers. The refined signal-to-sequence alignment approach was used to identify XNA signal slices that likely contain a NCB. To extract the XNA slices, we kmer-segmented the chunks employing DTW with our XNA kmer model. When selecting which XNA slices will be used to substitute the DNA signals, we randomly sampled five slices and then selected the one with the number of signals closest to the length of the insertion region. If the selected XNA slice was longer than the insertion region, we removed evenly spaced signal points as needed. Conversely, if the slice was shorter, we used linear interpolation to add evenly spaced signal points.

## Code and data availability

Source code for scripts used to obtain the XNA basecaller and analyze the data are available in a GitHub project at https://github.com/CSB5/XNA_Basecaller. Nanopore sequencing data is available from the European Nucleotide Archive (ENA – https://www.ebi.ac.uk/ena/browser/home) under project accession number PRJEB82716.

## Acknowledgements

This work was funded by the Advanced Manufacturing and Engineering (AME) Programmatic grant A18A9b0060. Additional funding supporting manpower came from Singapore Ministry of Health’s National Medical Research Council under its Open Fund – Individual Research Grants (NMRC/OFIRG/MOH-000649-00), and the Programme for Research in Epidemic Preparedness and Response (PREPARE), under its Joint Strategic Open Grant Call (Environmental Transmission & Mitigation and Diagnostics Co-operatives; PREPARE-OC-ETM-Dx-2023-006). Additional compute resources were provided by the A*STAR Computational Resource Center through the use of its high-performance computing facilities.

---

**Supplementary Figure 1.**
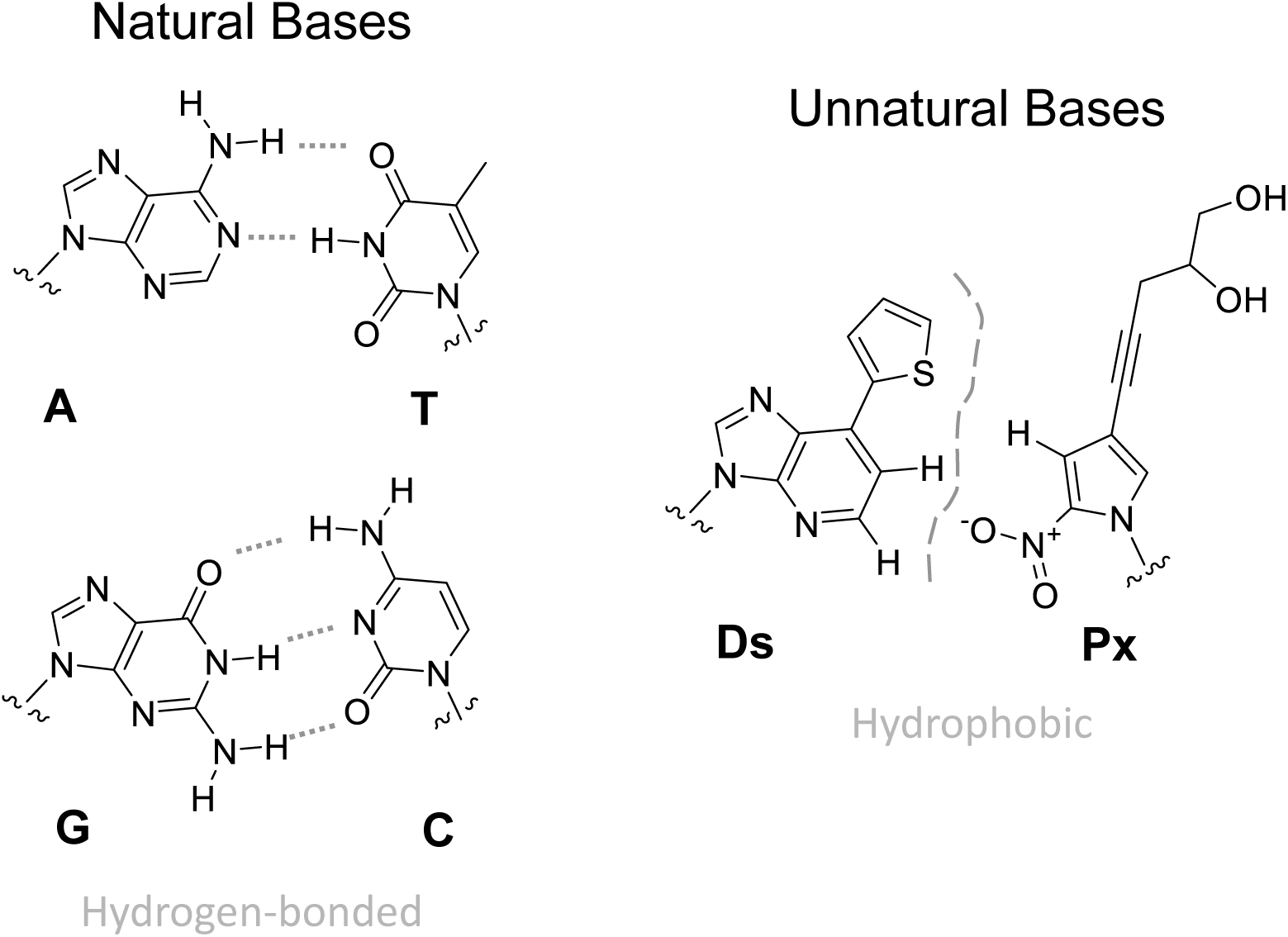
Chemical structures of natural and unnatural bases included in this study. The diagram shows how the natural (A with T, and G with C) and unnatural bases (Ds with Px) basepair. As can be seen here, the chemical structure of Ds and Px, and therefore of the corresponding XNAs containing them, can be quite distinct from natural DNA.

**Supplementary Figure 2.**
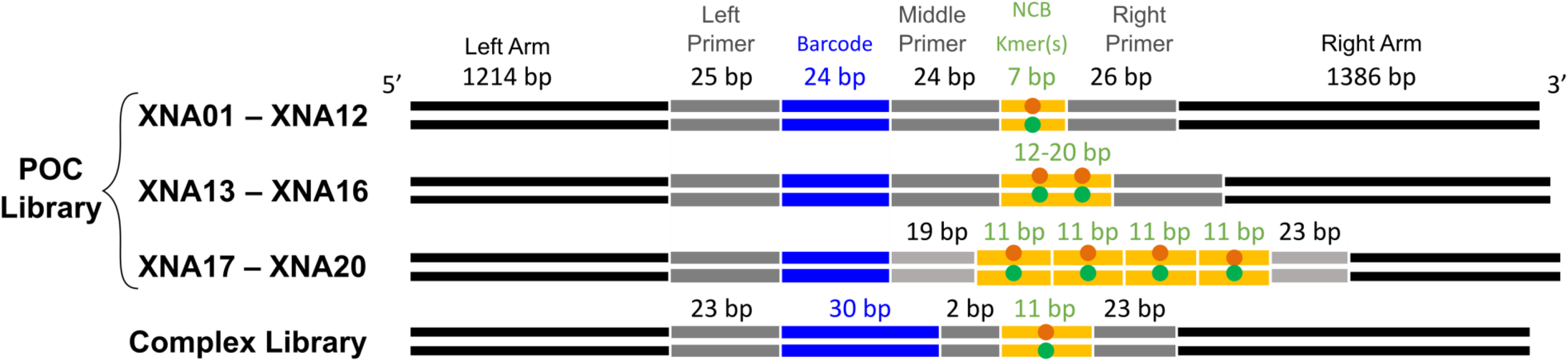
Design overview for XNA templates. Circles indicate the location of NCBs, Px in red and Ds in green respectively, on forward and reverse strands. All templates have the same left and right arm sequences. The primers (left, middle and right) are non-unique sequences surrounding the barcode and NCB kmer regions that are leveraged during synthesis (**Supplementary File 1** and **4**; **Methods**).

**Supplementary Figure 3.**
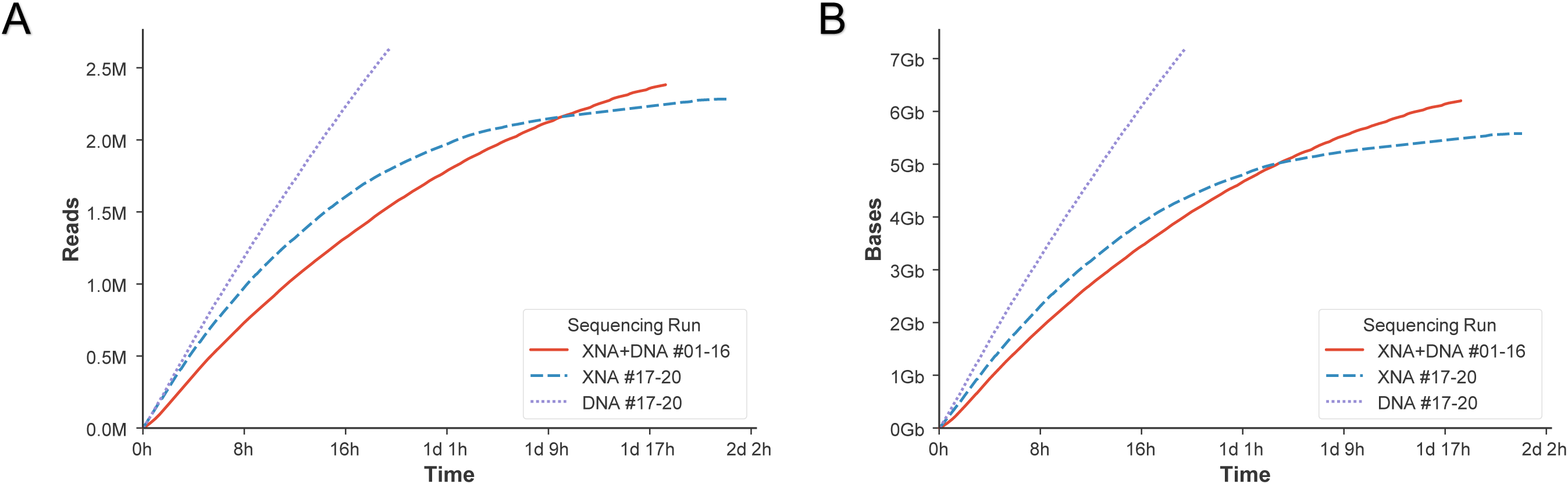
Proof-of-concept library sequencing throughput for XNA, DNA and mixed runs. The curves show cumulative sums for (A) number of reads and (B) bases sequenced as a function of time. XNA and DNA runs included templates 17-20, while the mixed run included XNA and DNA templates 1-16.

**Supplementary Figure 4.**
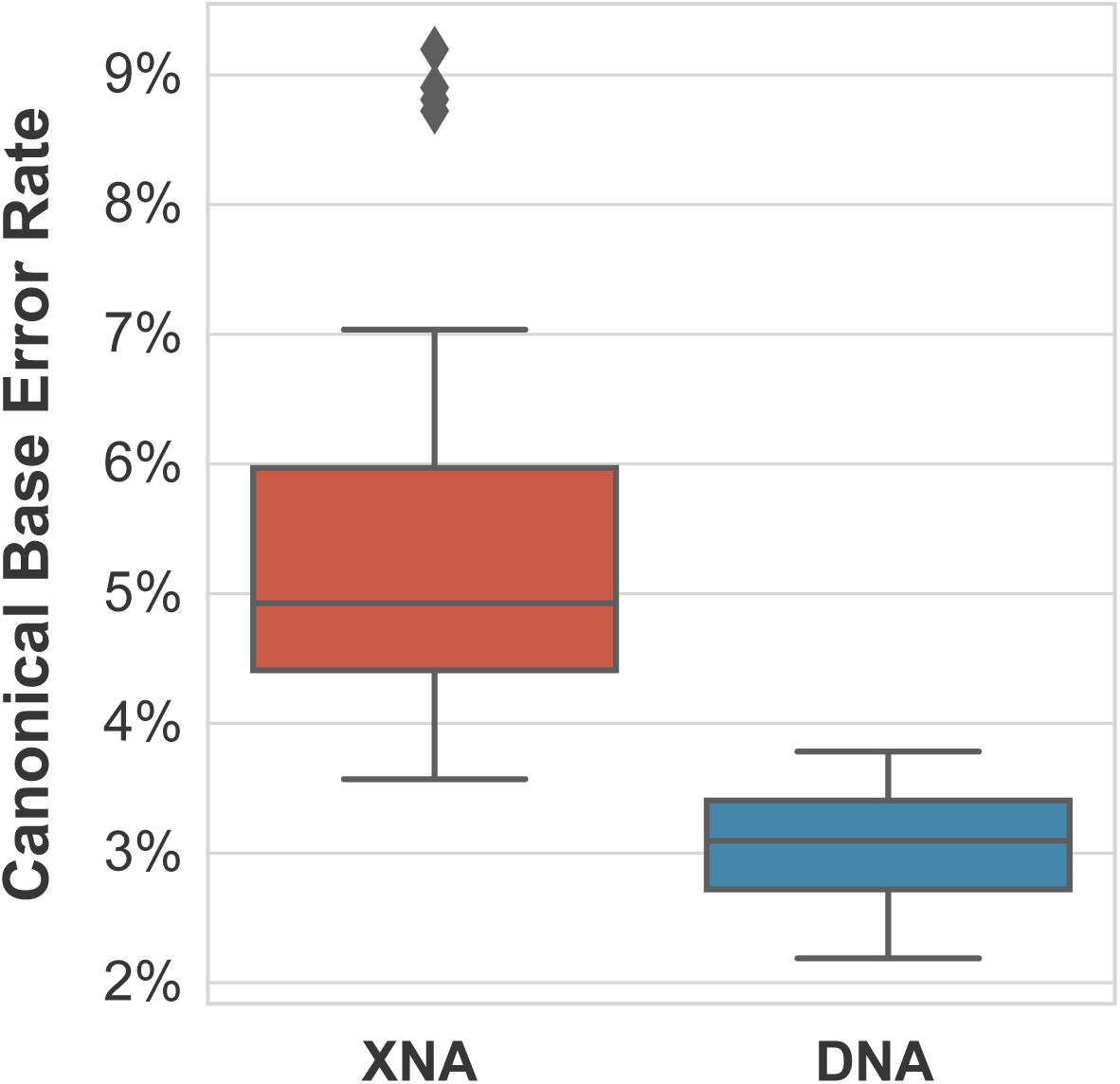
Basecalling error rates for canonical bases in XNA and DNA templates.

**Supplementary Figure 5.**
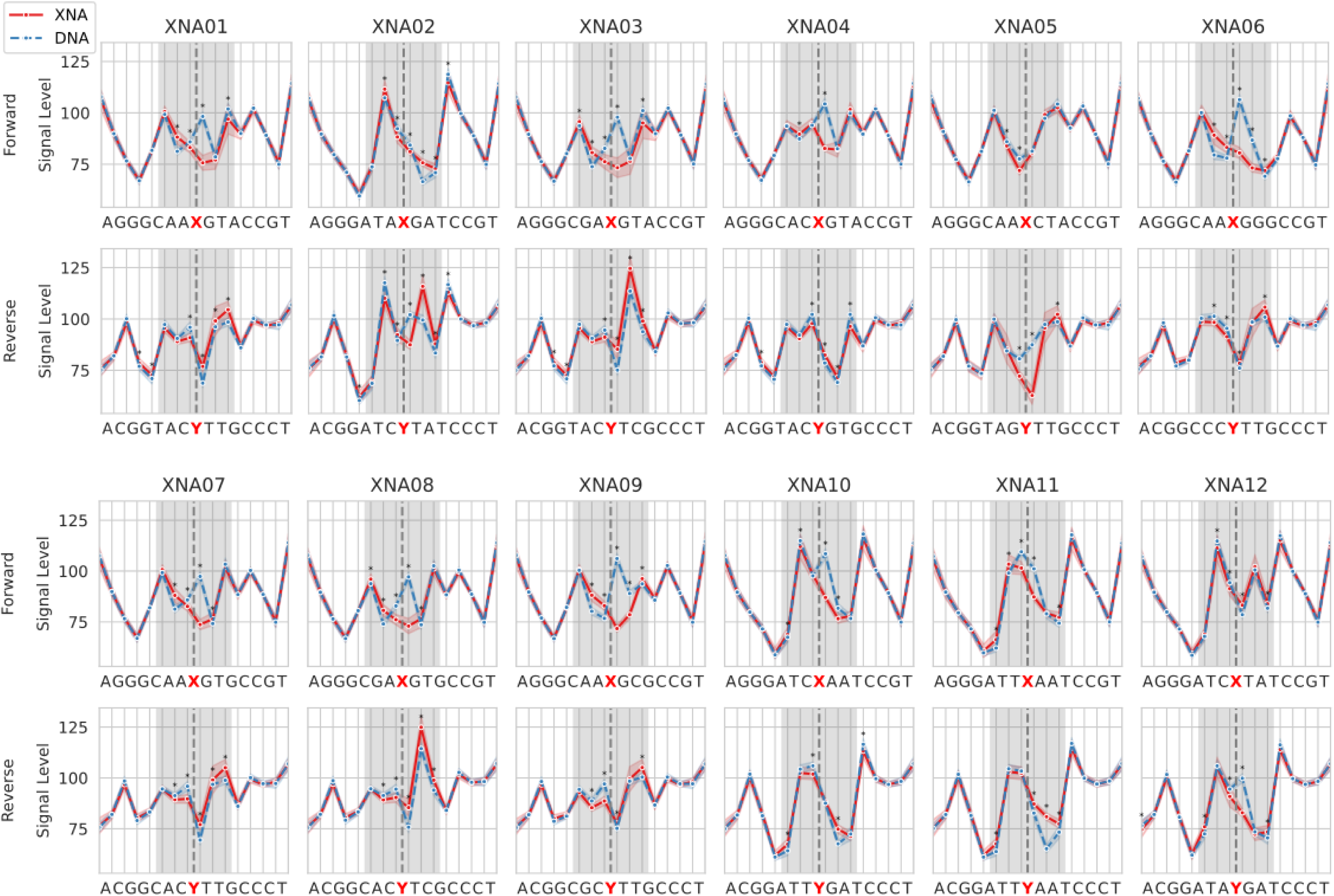
Nanopore electrical signal levels for XNA and DNA libraries around NCBs. Each curve shows average signal level based on a subsampling of 5,000 reads. Vertical dashed lines indicate the location of the NCB, shaded areas indicate a window of 3bp around the NCB, and asterisks (*) indicate Bonferroni adjusted two-sided Wilcoxon p-value<0.05 and signal fold-change>2%. All templates in this figure have a single non-canonical base in the XNA.

**Supplementary Figure 6.**
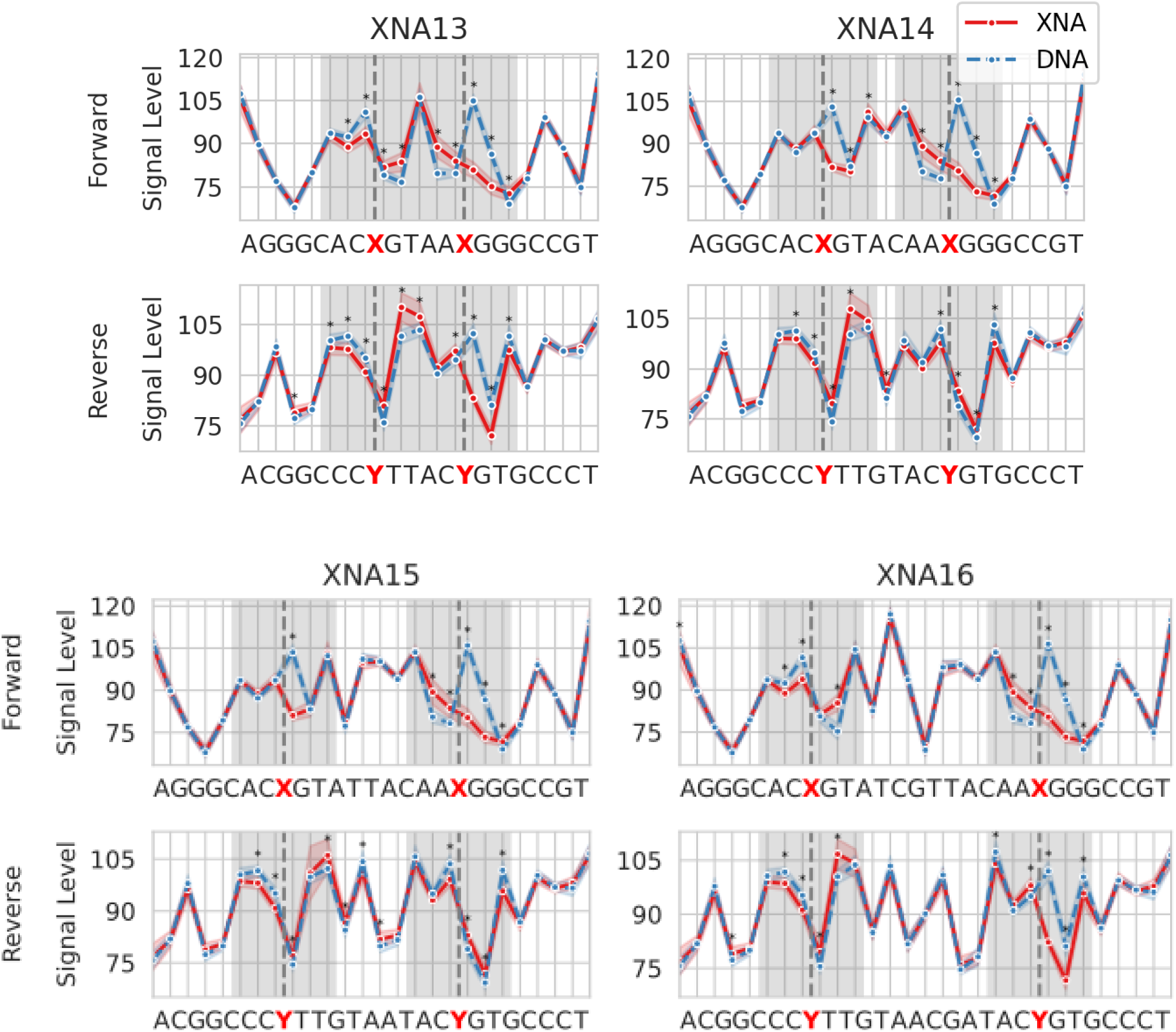
Nanopore electrical signal levels for XNA and DNA libraries around 2 consequent NCBs. Similar to **Supplementary Figure 5**, except that all templates in this figure have two non-canonical bases in the XNA.

**Supplementary Figure 7.**
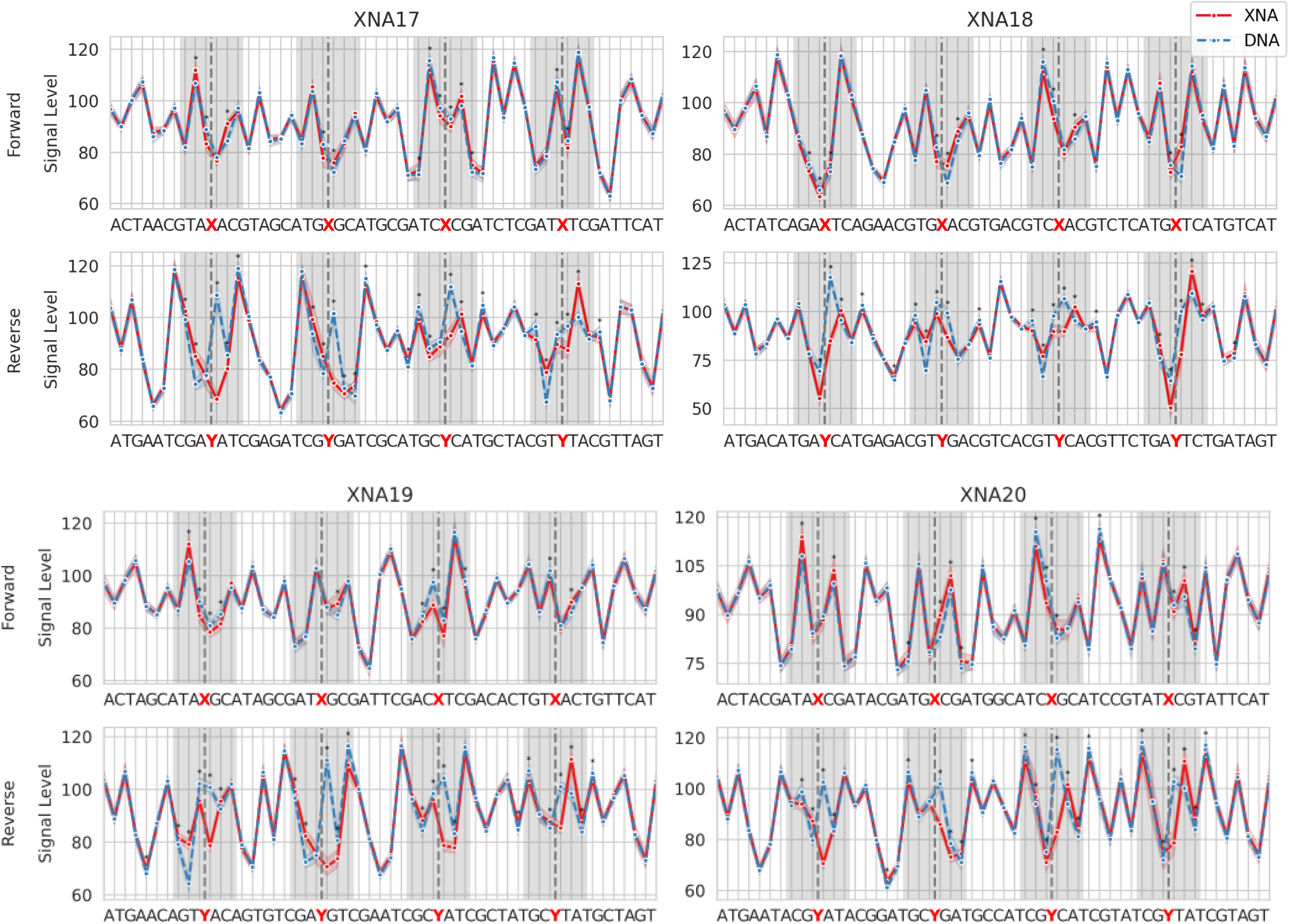
Nanopore electrical signal levels for XNA and DNA libraries around 4 consequent NCBs. Similar to **Supplementary Figure 5**, except that all templates in this figure have four non-canonical bases in the XNA.

**Supplementary Figure 8.**
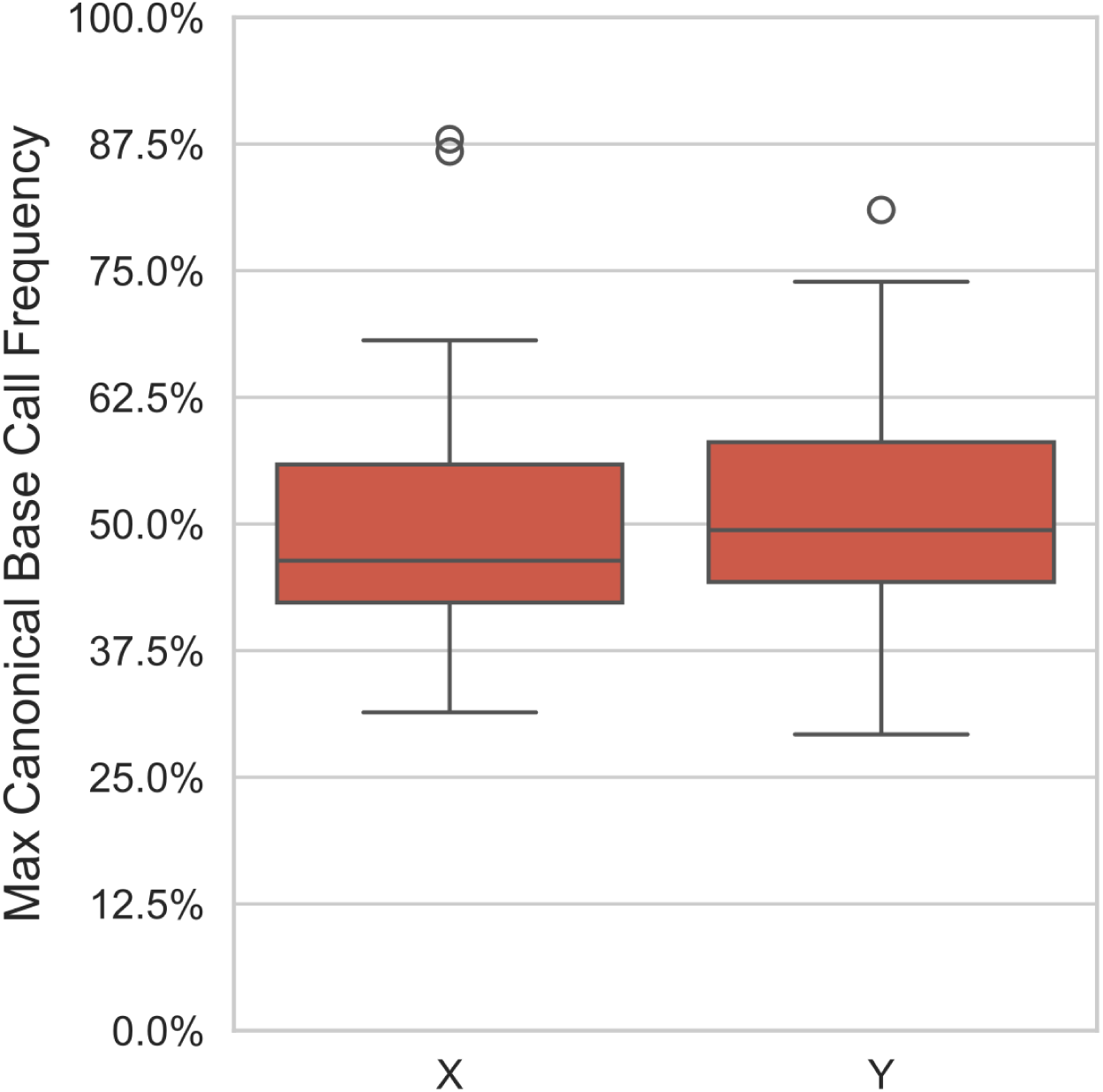
Native basecaller produces reproducible errors for non-canonical bases. Distribution of basecalling frequency of the most frequently called canonical base at an NCB position across templates in the proof-of-concept library.

**Supplementary Figure 9.**
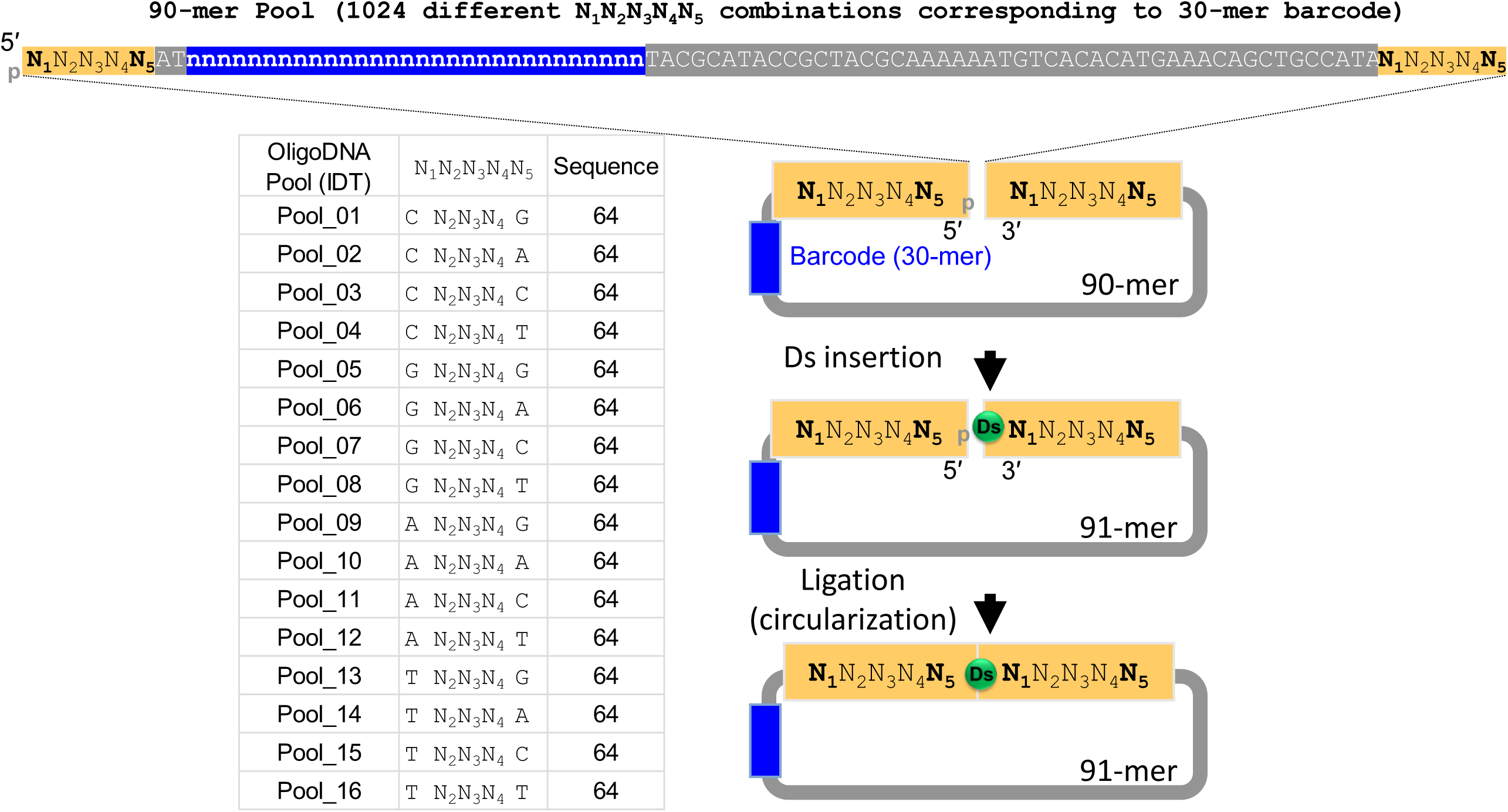
Schematic for Ds insertion process and corresponding oligo DNA pools generated. 16 different oligo DNA library pools, each of which consists of 64 defined fragments (50 pmole each, 90-mer) were generated.

**Supplementary Figure 10.**
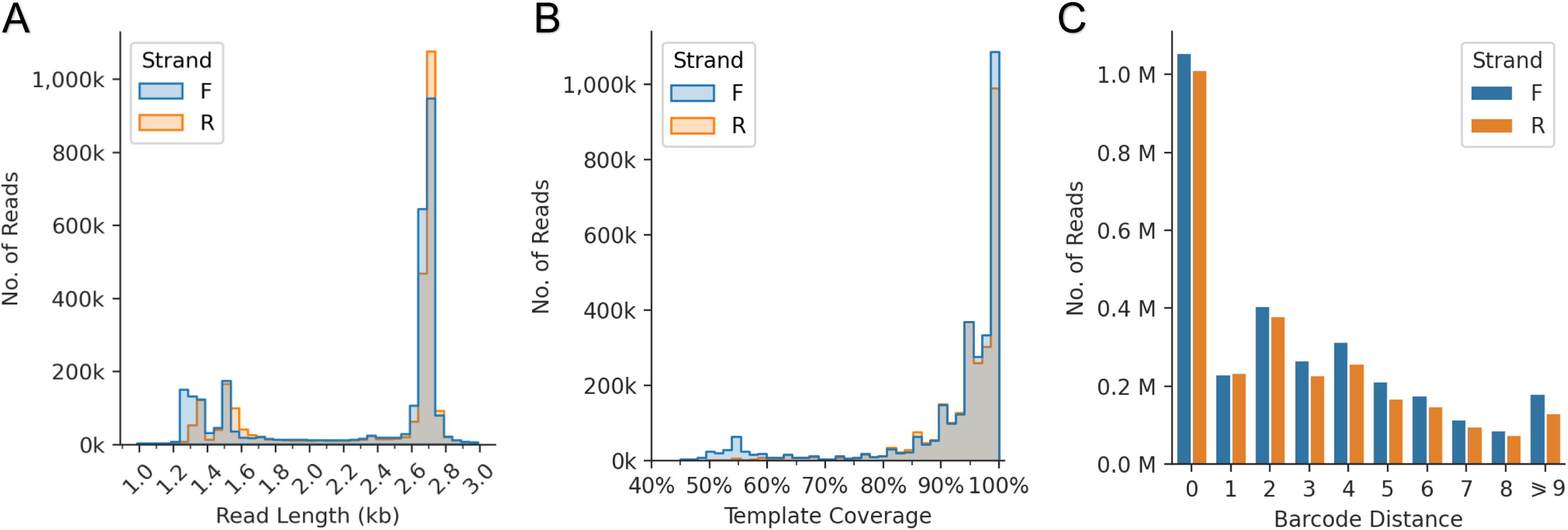
Sequencing statistics for the complex library. (A) Histogram of read lengths for forward and reverse strand reads. (B) Histogram of template coverage for template-specific region. (C) Bar chart with read counts as a function of barcode distance (computed as edit distance of sequenced barcode to the closest reference barcode).

**Supplementary Figure 11.**
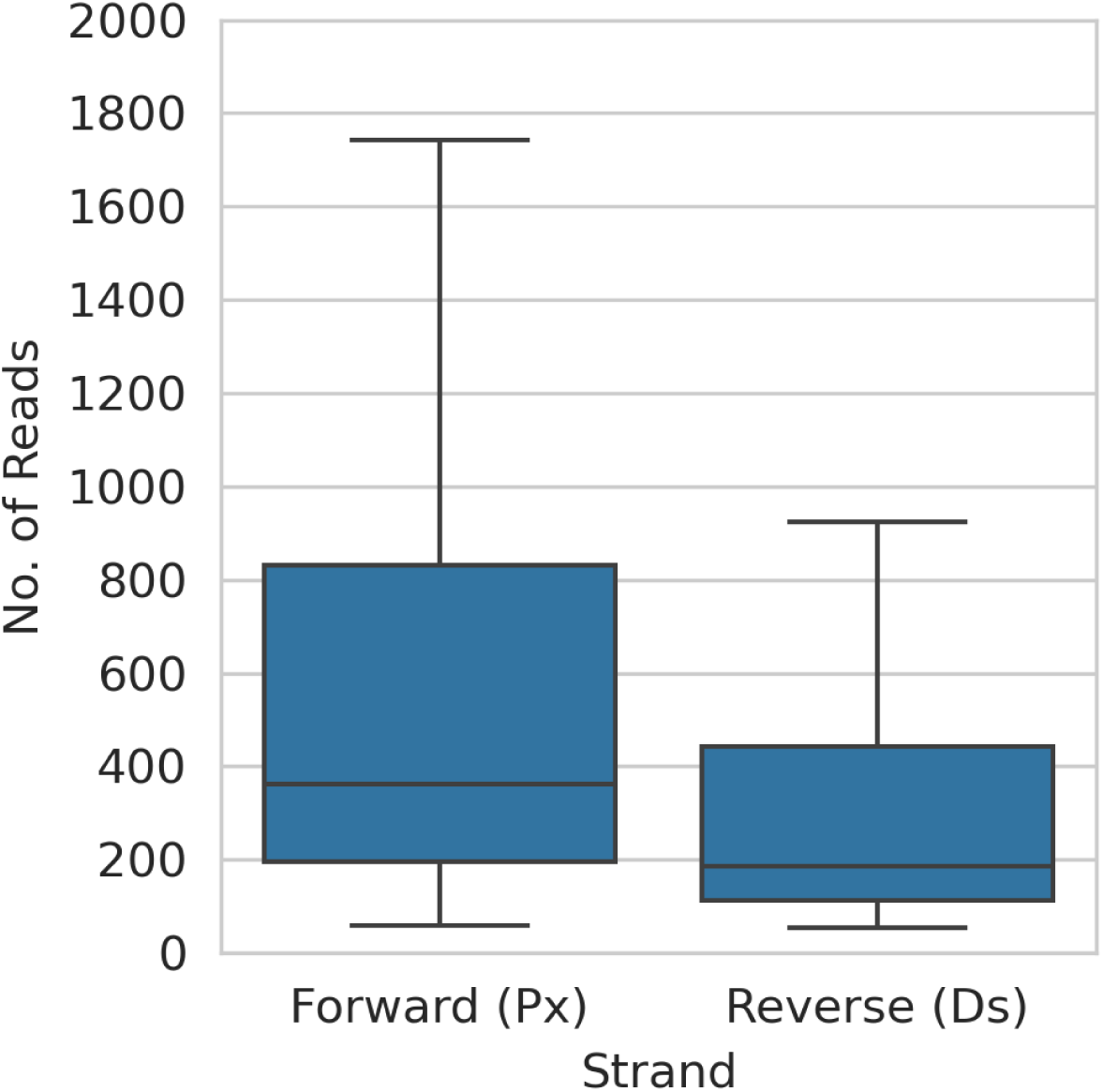
Read distribution per template in the complex library. Read counts for each template and strand for the two sequencing runs of the complex library, after filtering of reads by barcode distance (≤8), template coverage (≥85%) and read length (between 1k and 3k).

**Supplementary Figure 12.**
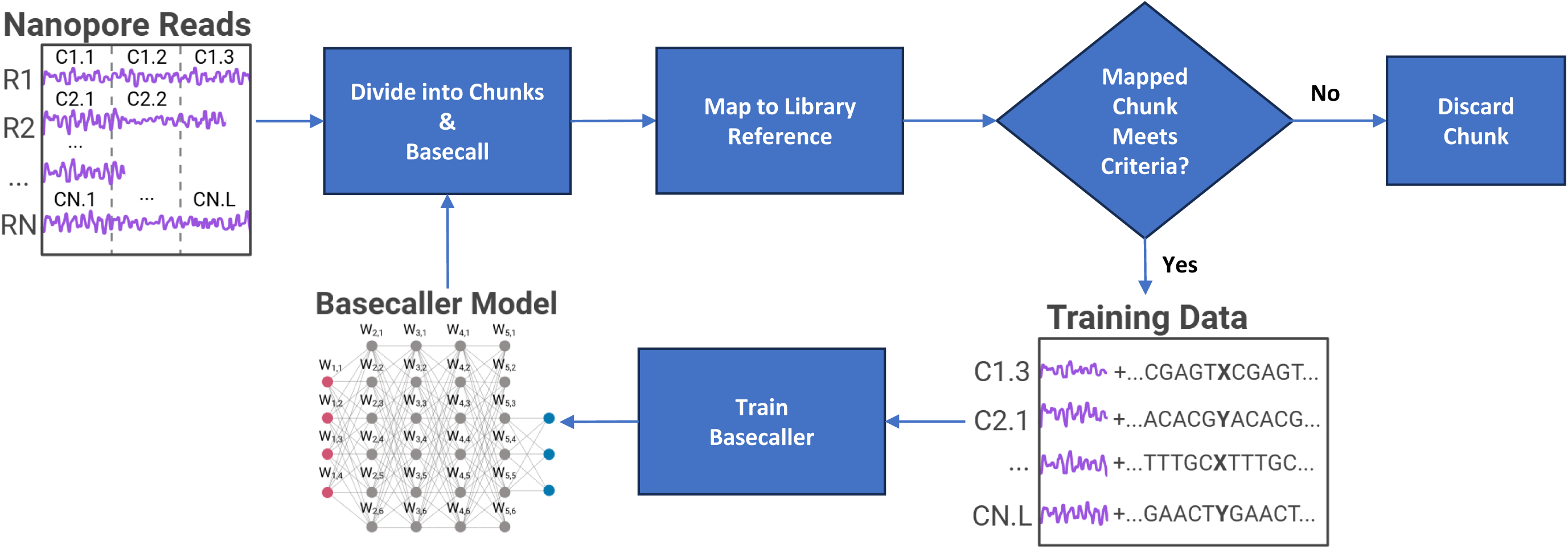
Bootstrapping methodology. At the beginning of each round of training, the signals from all Nanopore reads are divided into smaller chunks of signals, basecalled and mapped to the library reference (**Methods**). Mapped chunks with coverage and accuracy lower than 90% are discarded, and the remaining are added as training data for this round (paired signal chunks and reference sequences). The basecaller is trained using the acquired chunks, generating a new basecaller model. The same framework can now be repeated for the next round of training, using the basecaller model from the previous round to basecall the nanopore signal chunks. After each round of training more chunks meet the mapping criteria and can be used for training (**Supplementary Table 1**).

**Supplementary Figure 13.**
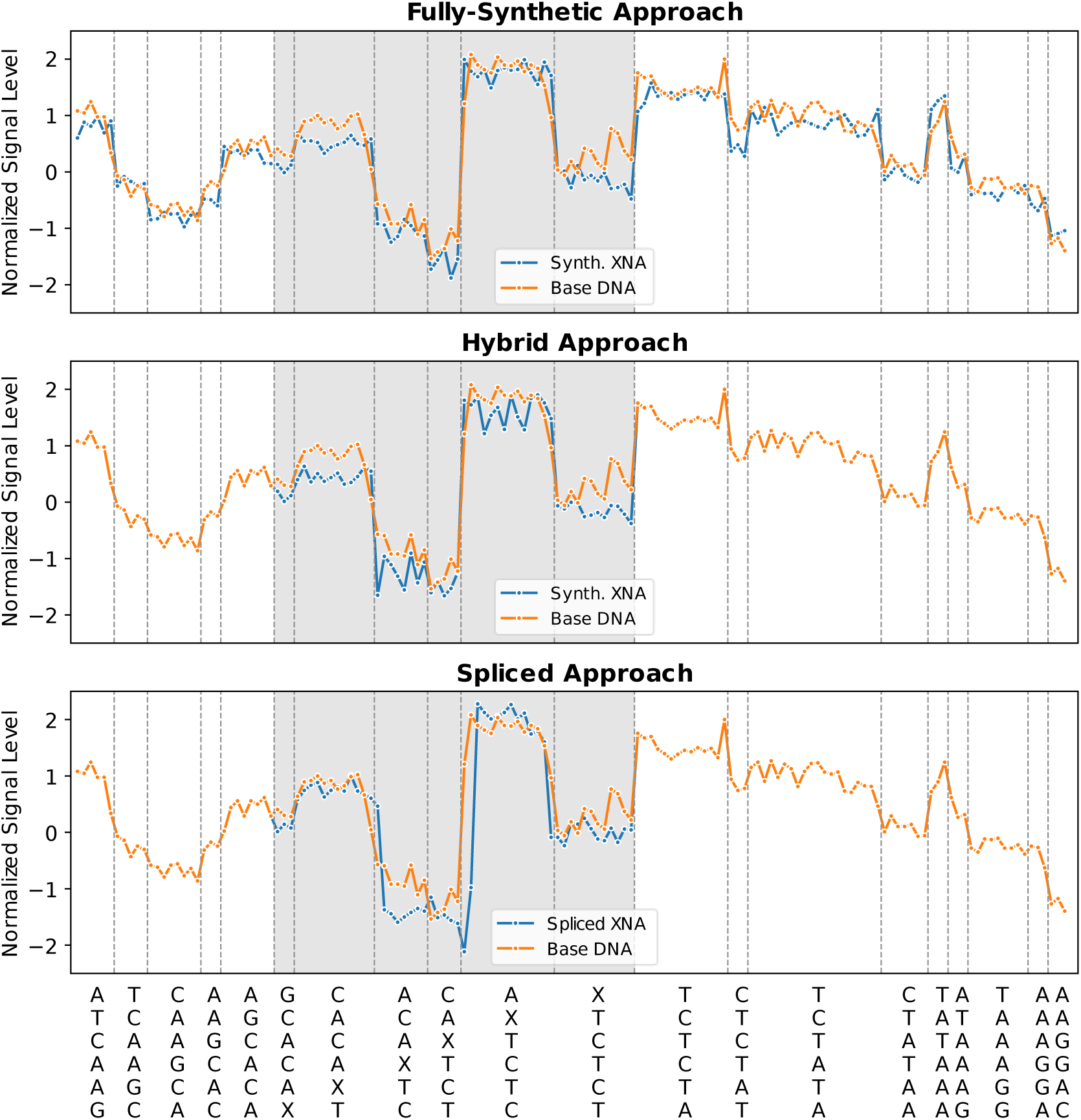
Training data examples per data augmentation approach. Each data point represents a nanopore measurement for the kmer indicated on x-axis. “Base DNA” and “Spliced XNA” refer to real signals, while “Synth. XNA” refers to simulated signals. Shaded areas indicate the signals of kmers containing an NCB. The base DNA sequence is “ATCAAGCACA**<u>G</u>**TCTCTATAAAGGAC”, where the bold and underlined **<u>G</u>** base is replaced by X.

**Supplementary Figure 14.**
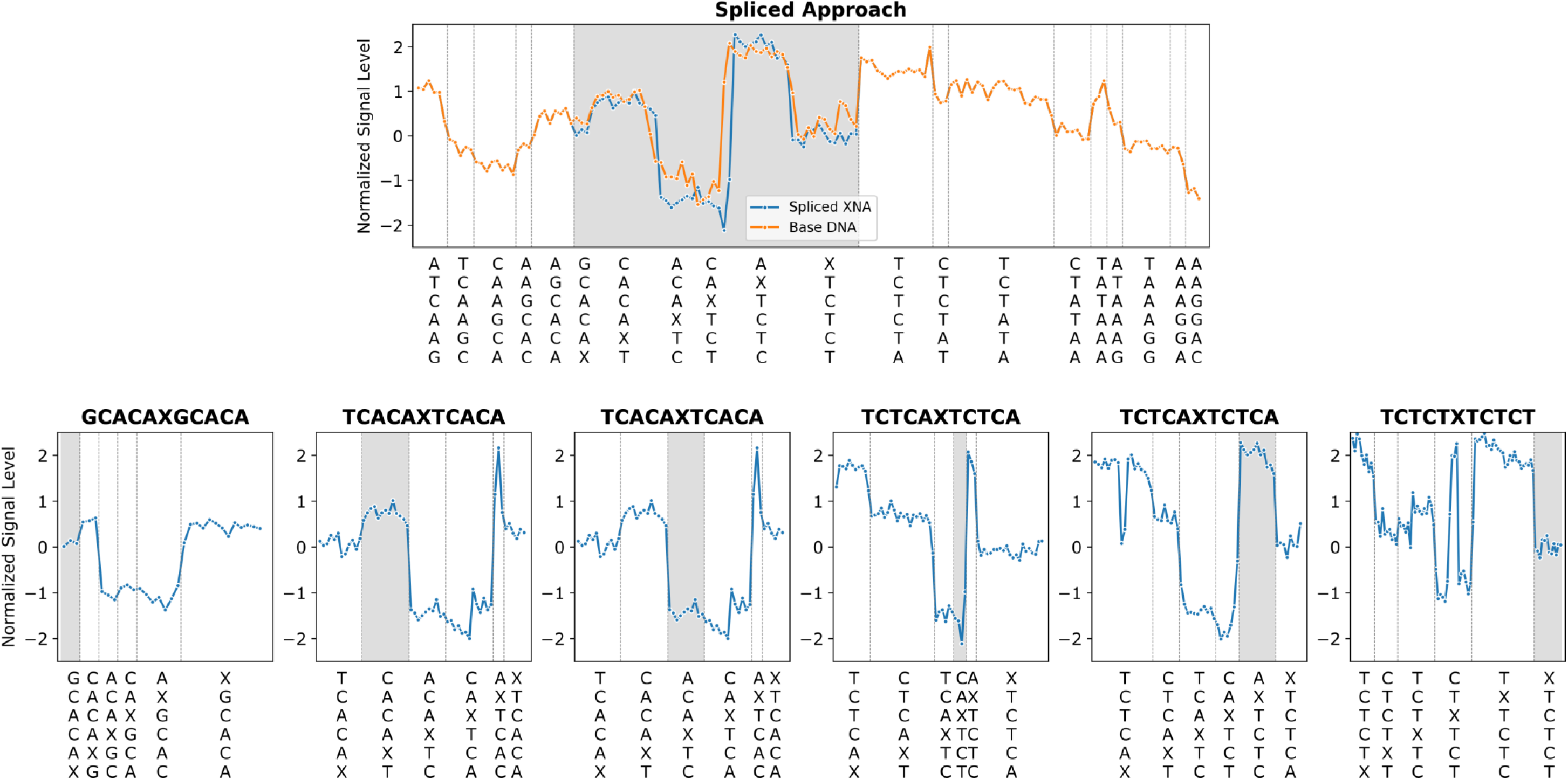
Spliced data-augmentation example. The newly formed spliced XNA chunk (shaded area in upper row figure) is assembled from slices coming from multiple real XNA chunks (shaded areas in lower row figures).

**Supplementary Figure 15.**
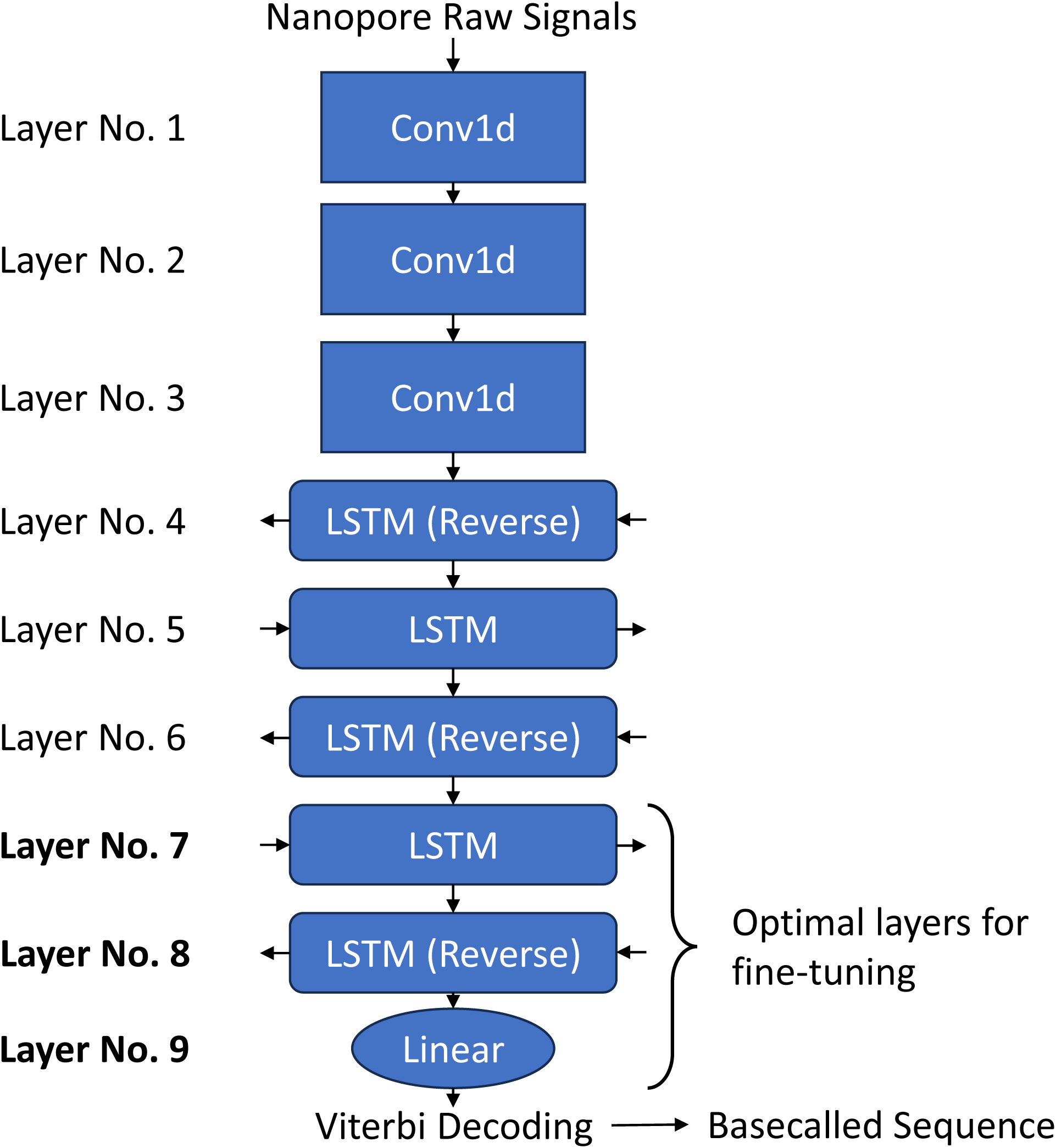
XNA basecaller architecture. The neural network architecture for the Bonito basecaller was suitably modified to enable basecalling of NCBs. Firstly, the output alphabet was expanded from four to six bases. Secondly, as the number of parameters for the last layer (layer No. 9, Linear) is exponentially related to the length of the alphabet and the chosen number of states, we reduced the number of states from 5 to 3 so that the model could fit into GPU memory for efficient training. Finally, the last 3 layers were fine-tuned to enable direct basecalling of XNAs.

**Supplementary Figure 16.**
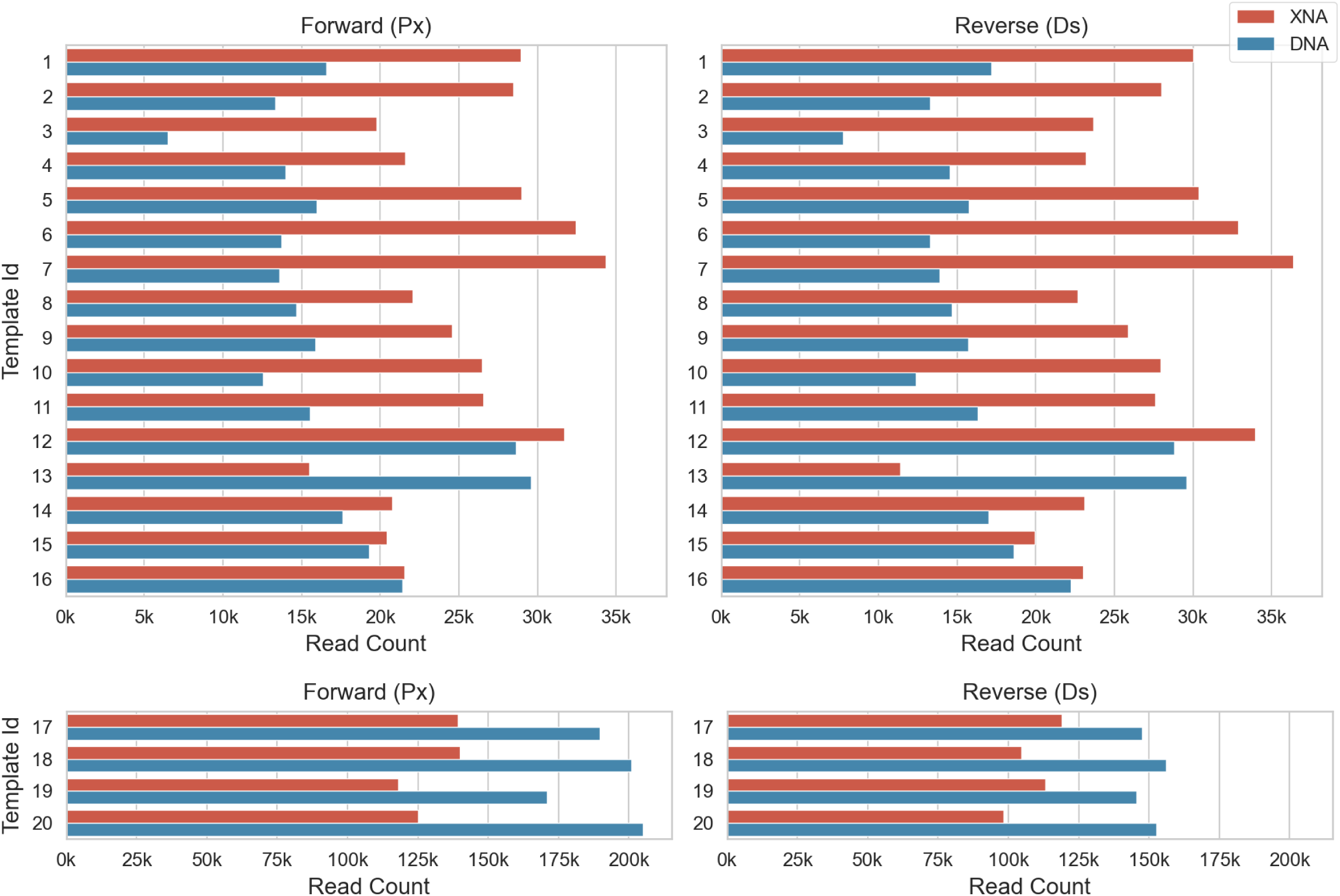
Read count per template for the proof-of-concept library. Reads were counted after selecting for those with appropriate barcode distance (≤5), template coverage (≥85%) and read length (between 1k and 3k).

**Supplementary Table 1.**
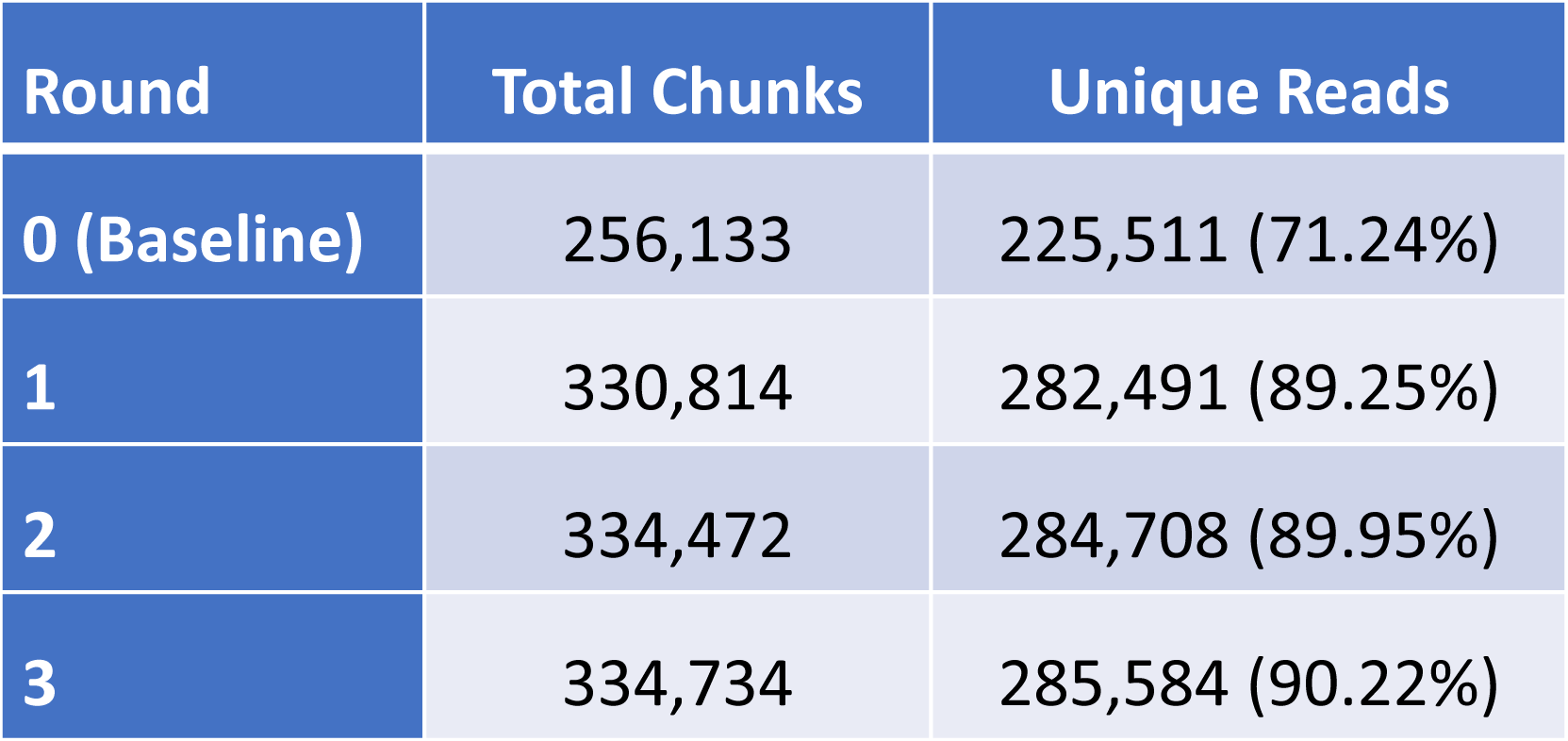
XNA train data signal chunks summary. . Summary of the number of signal chunks satisfying the filter criteria (>90% coverage and accuracy), obtained from basecalling train and validation set XNA reads with the bootstrapped model from different rounds of training.

| Round | Total Chunks | Unique Reads |
| --- | --- | --- |
| 0 (Baseline) | 256,133 | 225,511 (71.24%) |
| 1 | 330,814 | 282,491 (89.25%) |
| 2 | 334,472 | 284,708 (89.95%) |
| 3 | 334,734 | 285,584 (90.22%) |

